# Piezo1 balances focal and reticular adhesions to enable EGFR clathrin-mediated endocytosis

**DOI:** 10.64898/2026.08.21.746250

**Authors:** Oriol Bagudanch, Ona Lupiáñez, Valentina Ayala, Roy Midyan, Dustin C. Bagley, Satish Babu Moparthi, Zaber Shabib, Laura Hakanpää, An-Sofie Lenaerts, Leonardo Almeida-Souza, Francisco J. Muñoz, Miguel A. Valverde, Stéphane Vassilopoulos, Jody Rosenblatt, Carlos Pardo-Pastor

**Affiliations:** Laboratory of Molecular Physiology, Department of Medicine and Life Sciences, Universitat Pompeu Fabra, Barcelona, Spain; Randall Centre for Cell and Molecular Biophysics, New Hunt’s House, School of Basic and Medical Sciences, Faculty of Life Sciences and Medicine, King’s College London, London, United Kingdom; The Francis Crick Institute, London, United Kingdom; Sorbonne Université, INSERM, Institute of Myology, Centre of Research in Myology, UMRS 974, Paris, France; Helsinki Institute of Life Science, Institute of Biotechnology and Faculty of Biological and Environmental Sciences, University of Helsinki, Helsinki, Finland; School of Cancer and Pharmaceutical Sciences, King’s College London, London, United Kingdom

**Keywords:** Piezo1, RhoA, focal adhesion, clathrin, endocytosis, reticular adhesion, flat clathrin lattice, clathrin plaque, Yoda1, Fyn, VAV2, N-WASP, Arp2/3, EGFR

## Abstract

Cells attach to the extracellular matrix through distinct integrin-mediated adhesive structures, including force-transmitting focal adhesions (FAs) and clathrin-enriched reticular adhesions (RAs). FAs enable mesenchymal cell migration and disassemble at mitotic entry, whereas RAs impede migration, persist during mitosis, and contribute to clathrin-mediated endocytosis (CME) as they disassemble. FAs grow with RhoA contractility, whereas RAs shrink, but the mechanisms coordinating these opposing responses remain unclear. Here, we identify the mechanically activated ion channel Piezo1 as a master regulator of FA/RA balance. Piezo1-dependent calcium influx activates the Src family kinase Fyn, which activates two actin polymerization pathways: FA and stress fiber growth via VAV2-RhoA and RA disassembly via N-WASP-Arp2/3. Inhibition or knockdown of Piezo1, Fyn, or VAV2 decreases FA size and increases RA coverage. Critically, cells lacking Piezo1 fail to internalize ligand-activated EGFR on stiff substrates despite normal CME on soft substrates, establishing an essential role for Piezo1 in EGFR CME mechanoadaption. Our findings reveal Piezo1 as the mechanosensor linking membrane tension to coordinated actin polymerization pathways that co-regulate cell-matrix adhesion and endocytosis. Given that CME contributes to viral entry into host cells and cancer resistance to anti-EGFR antibody therapy, targeting the Piezo1-RA-CME axis may offer novel therapeutic opportunities.

## Introduction

In solid tissues, cell attachment to the surrounding extracellular matrix (ECM) is essential for fundamental processes including cell cycle progression, mitosis, differentiation, and migration. Integrin-mediated adhesions link the ECM across the plasma membrane to the cell cytoskeleton and serve as biochemical and mechanical signaling hubs^1^. Among these, focal adhesions (FAs) transmit actomyosin-generated forces and grow in response to RhoA GTPase activation by both soluble and mechanical extracellular signals^2,3^. Cells also attach to the ECM via reticular adhesions (RAs), which contain flat clathrin lattices (also called clathrin plaques), require β5 integrin, and link to intermediate filaments and Arp2/3-dependent branched actin rather than to the RhoA-dependent actomyosin cytoskeleton of FAs^1,4–9^. Beyond differences in composition, FAs and RAs have opposing functions: FAs transmit actomyosin forces to the ECM, enabling mesenchymal migration and YAP nuclear accumulation, whereas RAs are abundant in static cells (e.g., myocytes), serve as extranuclear YAP anchors, and must disassemble to allow migration^2,6,10–12^. FAs and RAs also show opposing behaviors: in interphase cells, FAs grow with RhoA-dependent actomyosin contractility, whereas RAs shrink, and upon mitotic entry, FAs disassemble, whereas RAs persist as essential mitotic adhesions, with their perturbation causing multinucleation, frequent in cancer^4^. Yet, these opposing mechanoresponses remain poorly understood.

Crucially, RA disassembly gives rise to clathrin-coated vesicles, contributing to clathrin-mediated endocytosis (CME), the main membrane internalization pathway in eukaryotic cells^9,13,14^. Flat clathrin lattices and their partial disassembly into CME vesicles were first described by electron microscopy studies almost five decades ago^9,15,16^. Yet, the forces driving clathrin rearrangement and membrane deformation during RA disassembly remained largely unknown until recent work identified N-WASP-Arp2/3-mediated branched actin polymerization as essential for RA disassembly and CME of specific cargoes like epidermal growth factor receptor (EGFR)^5,14,17^.

By promoting cell-substrate attachment while suppressing CME, RAs may transiently isolate mitotic cells from environmental signals, contributing to mitotic robustness, and FA-RA toggling in interphase could tune mechanical and chemical signaling without requiring additional receptors^4,12,14,18^. What controls this toggling remains unclear, but membrane mechanical tension is an excellent candidate: it increases during migration, opposes the membrane deformation required for CME vesicle formation, and in mechanically challenging conditions (hypotonicity, stiff substrates), CME is maintained by N-WASP-Arp2/3-mediated branched actin polymerization^5,14,19–23^. However, how membrane tension activates this CME switch and coordinates opposing FA and RA responses remains unclear. Given that the ion channel Piezo1 is activated by membrane tension and promotes both FA growth and CME^24–29^, we hypothesized that it could also couple membrane tension increases to Arp2/3-dependent RA disassembly and CME. Here we reveal that Piezo1 simultaneously promotes FA growth via VAV2-RhoA and RA disassembly via N-WASP-Arp2/3, identifying it as the mechanosensor that couples membrane tension to coordinated actin polymerization pathways that co-adapt adhesion and endocytosis to the mechanical conditions of the cell environment^5,12,14,20,21^.

## Results

### Piezo1 balances FAs and RAs via RhoA

To directly test if Piezo1 affects FAs and RAs, we treated serum-starved HeLa cells with the Piezo1 agonist Yoda1^30^ and immunostained them for β5 integrin and the FA marker Paxillin. Given that FAs and RAs share β5 integrin but only FAs contain Paxillin, we scored (β5 integrin^+^/Paxillin^+^) as FAs and (β5 integrin^+^/Paxillin^-^) structures as RAs^4,10^. Yoda1 treatment produced longer FAs and more stress fibers, while simultaneously decreasing RAs, quantified as the percentage of cell area covered by RAs^10^ (Fig. 1a-c). Similar results were obtained with indirect RhoA activation by microtubule depolymerization with nocodazole^31^ or irreversible RhoA activation by Rho Activator II (a cell-permeable recombinant bacterial toxin) (Fig. S1a-b). In contrast, Rho inhibition with cell-permeable C3 transferase reduced FA size and increased RA coverage, as expected^3,12,32^ (Fig. 1c, f). Importantly, Rho inhibition suppressed Yoda1 effects (Fig. 1d-f). Therefore, Piezo1 activation shifts RAs to FAs via RhoA GTPase. Moreover, these experiments suggest that Piezo1 balances FAs and RAs, as FAs growth is matched to RAs loss.

**Figure 1.**
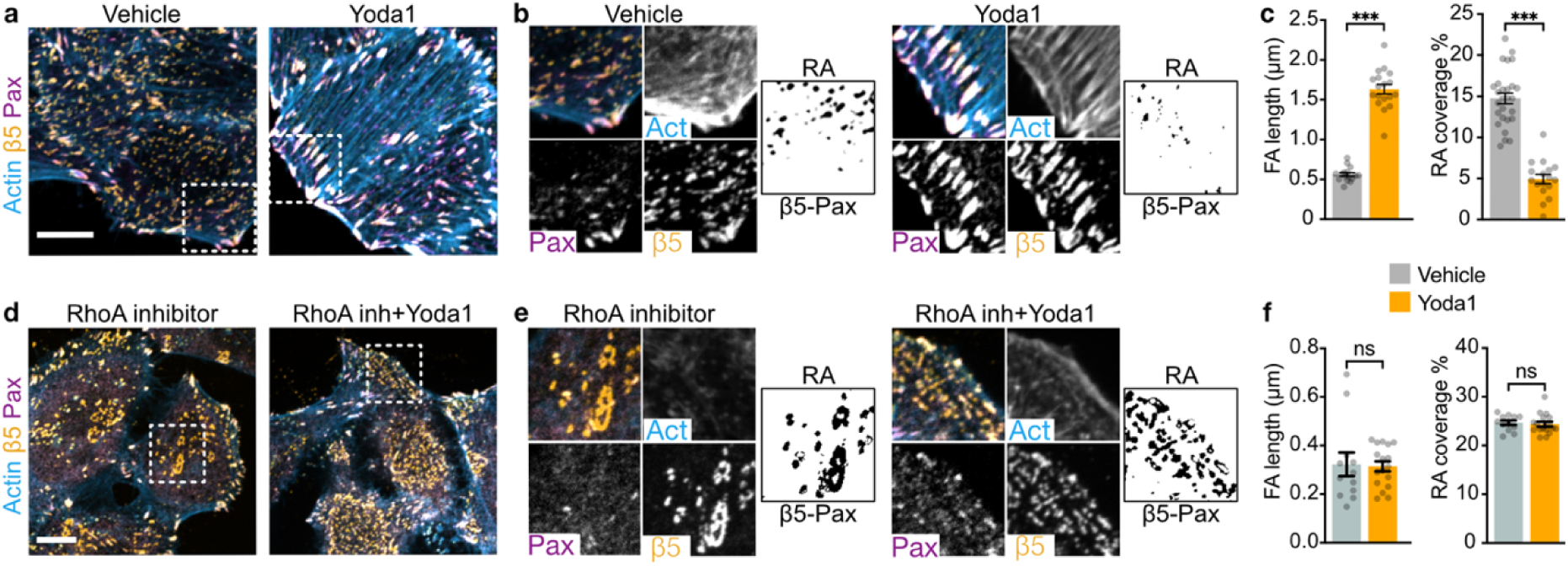
Piezo1 balances FAs and RAs via RhoA. **a.** Representative confocal images of HeLa cells stained as indicated after overnight serum starvation and 2h treatments with DMSO (vehicle control) or 10 µM Yoda1. White squares define the regions zoomed in b. Scale bar: 10μm. **b.** Zooms from a. RA image is the result of subtracting β5-Pax signals (for details, see materials and methods). **c.** Bar plots representing mean±SEM FA length from Pax stainings (left) or RA coverage expressed as the cell are percentage occupied by RA (right) in cells treated as indicated. N: Vehicle=16, Yoda1=18 cells from 3 experiments. **d.** Representative confocal images of HeLa cells stained as indicated after 2h treatments with DMSO (vehicle control) or 10 µM Yoda1 in the presence of 0.5μg/mL Rho Inhibitor (added 30min earlier). White squares define the regions zoomed in e. Scale bar: 10μm. **e.** Zooms from d. RA image is the result of subtracting β5-Pax signals (for details, see materials and methods). **f.** Bar plots representing mean±SEM FA length from Pax stainings (left) or RA coverage expressed as the cell are percentage occupied by RA (right) in cells treated as indicated. N: Rho inhibitor=12, Rho inhibitor+Yoda1=17 cells from 3 experiments. ns, non-significant; *** p<0.001; in Mann-Whitney test.

### Piezo1 knockdown promotes RAs

To test if impairing Piezo1 activity would shift FAs to RAs, we performed our FA/RA analysis in cells transiently transfected with Piezo1-targeting siRNA. Piezo1 knockdown (confirmed by suppression of Yoda1-induced calcium signals, Fig. S2a-c), reduced FAs and increased Ras (Fig. 2a-c), demonstrating that Piezo1 balances FAs and RAs. Acute Piezo1 inhibition with GsMTx4^33^ showed similar tendencies to shrink FAs and enlarge RAs (Fig. S2d-f), suggesting that Piezo1 activity, rather than indirect long-term changes due to Piezo1 knockdown, control FA/RA balance. Importantly, Piezo1 knockdown and inhibition suppressed Yoda1-, but not nocodazole-induced FA lengthening and RA loss (Fig. 2a-c, S2d-f), further confirming that RhoA acts downstream of Piezo1 and ruling out a general defect in RhoA-dependent FA formation and RA loss upon impairing Piezo1 expression or function.

**Figure 2.**
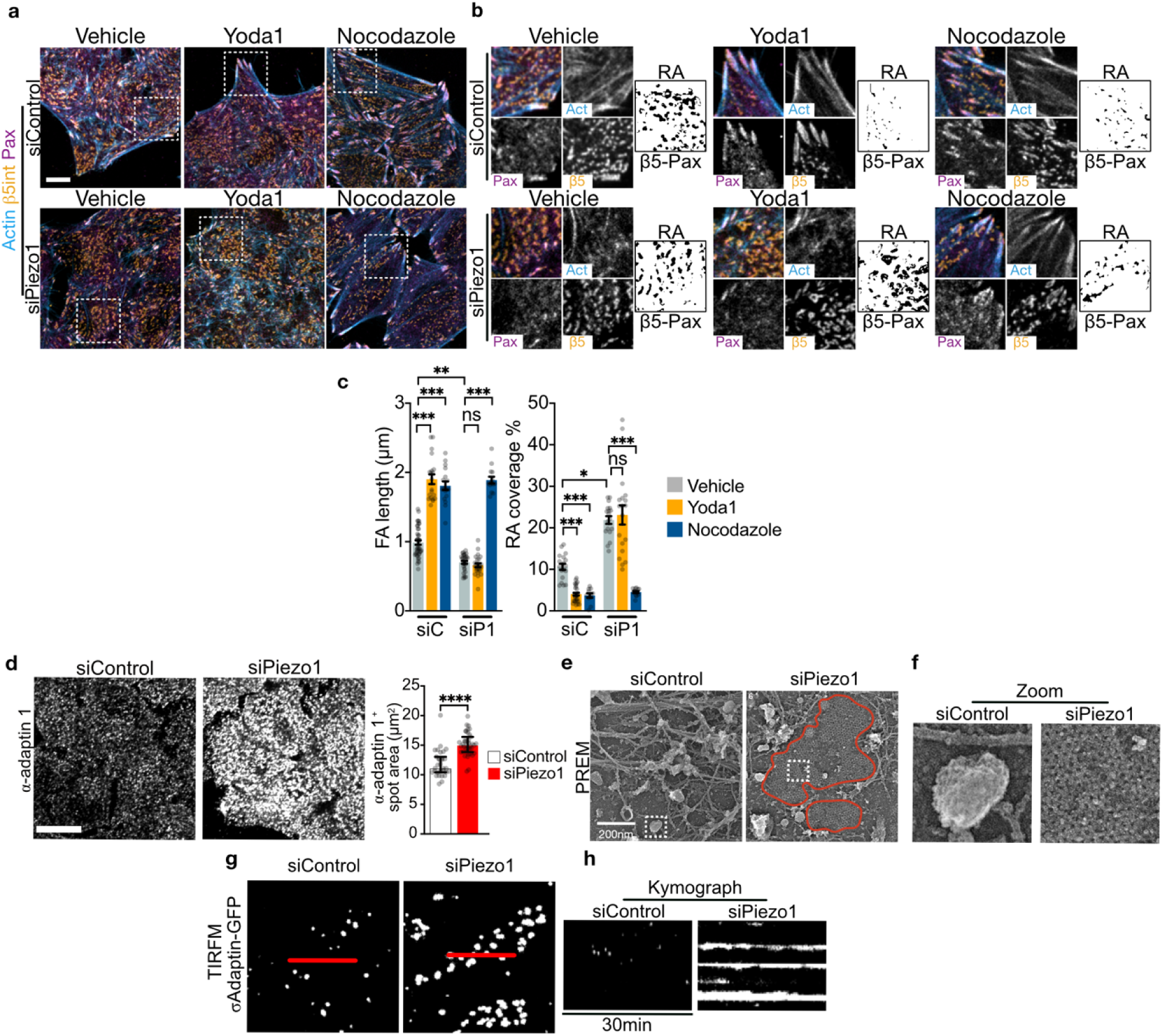
Piezo1 knockdown promotes RAs. **a.** Representative confocal images of HeLa cells transfected with siRNA for 48h, serum-starved overnight, treated for 2 hours with DMSO (vehicle control), 10 µM Yoda1, or 25µM Nocodazole (positive control), and stained as indicated. White squares define the regions zoomed in h. Scale bar: 10μm. **b.** Zooms from a. RA image is the result of subtracting β5-Pax signals (for details, see materials and methods). **c.** Bar plots representing mean±SEM FA length from Pax stainings (left) or RA coverage expressed as the cell are percentage occupied by RA (right) in cells treated as indicated. N, FA: Vehicle=43 (siControl) 30 (siPiezo1), Yoda1=20 (siControl) 24 (siPiezo1), Nocodazole=17 (siControl) 14 (siPiezo1) cells, RA: Vehicle=18 (siControl) 18 (siPiezo1), Yoda1=28 (siControl) 19 (siPiezo1), Nocodazole=11 (siControl) 13 (siPiezo1) cells from 3 siRNA transfections. **d.** Representative confocal images of α-Adaptin 1 stainings in HeLa cells serum-starved overnight after 48h-transfections with siRNA. N: siControl=31; siPiezo1= 37 cells from 3 siRNA transfections. **e.** Platinum-replica electron microscopy (PREM) images of siRNA-transfected cells. The red line surrounds flat clathrin structures corresponding to RA. White squares highlight clathrin-coated endocytic vesicles (siControl) and an RA portion with visible flat clathrin (siPiezo1), zoomed in f. Cells were transfected with siRNA for 72h. Scale bar: 200nm. **f.** Zooms from e. **g.** Stills of total interference reflection microscopy (TIRFM) videos of CRISPR-engineered U2OS cells expressing endogenous σAdaptin-GFP. Red lines were used to build kymographs shown in h. Cells were transfected with siRNA for 72h. **h**. Kymographs showing the evolution of endogenous σAdaptin-GFP signal along the red lines in m in siRNA-transfected cells. ns, non-significant; *p<0.05; **, p<0.01; *** p<0.001; **** p<0.0001 in Mann-Whitney (d), or ANOVA followed by Kruskal-Wallis with Dunn’s correction (c).

To further confirm that Piezo1 knockdown promotes RAs, we first immunostained siRNA-treated cells with additional RA components. α-adaptin 1 is a member of the AP-2 adaptor complex linking the plasma membrane to clathrin. Its immunostaining acts as a surrogate of clathrin dynamics, showing bigger and brighter spots at RA than at endocytically-active clathrin-coated pits^8,10,12,21^. Accordingly, α-adaptin 1 stainings revealed larger spots at the basal membrane of Piezo1-knockdown cells, compared to cells transfected with control siRNA (Fig. 2d). We next used platinum-replica electron microscopy (PREM), which reveals nanometer resolution of native clathrin structures in cells^7,9,12,17,34^, on adherent siRNA-treated cells exposed to an ultrasonic pulse that removes most cellular structures while preserving the ventral plasma membrane^35^. Piezo1 knockdown increased the area of flat clathrin structures, associated with RAs, at the ventral plasma membrane (Fig. 2e, f). Moreover, total internal reflection fluorescence microscopy (TIRFM) of RA dynamics in live cells^10,29^ revealed abundant, large σ-adaptin^+^ structures in the ventral membrane of Piezo1-knockdown cells (Fig. 2g). Kymograph analysis of these movies revealed longer-lived σ-adaptin^+^ structures in Piezo1-knockdown cells, a key feature of RAs^4,8–10^ (Fig. 2h). To avoid artifacts from overexpression, we relied on CRISPR-engineered U2OS cells expressing endogenous σ-adaptin fluorescently labelled, previously used in RA and CME studies^10,36^. Together, our findings confirm that Piezo1 inhibition and knockdown promote RAs at the expense of FAs, further supporting that Piezo1 balances FAs and RAs.

FA/RA balance by Piezo1 requires extracellular Ca^2+^

We next tested if Piezo1-RhoA-dependent FA/RA balance requires extracellular Ca^2+^, a ubiquitous second messenger integral to Piezo1/2-RhoA-adhesion dynamics^29,37–39^. Extracellular Ca^2+^ removal reduced FA length and actin fibers while increasing RA coverage and, importantly, prevented Yoda1-induced FA/RA changes (Fig. 3a-c), suggesting that both basal FA/RA balance and Piezo1-induced FA growth and RA shrinkage require extracellular Ca^2+^. However, nonspecific extracellular Ca^2+^ entry induced by the Ca^2+^ ionophore A23187 did not reproduce Yoda1-effects on adhesions (Fig. 3a, c, right), suggesting that Ca^2+^ entry is not sufficient to regulate FA/RA balance. Together, these results show that both baseline and Yoda1-induced FA/RA balance require extracellular Ca^2+^ influx via Piezo1.

**Figure 3.**
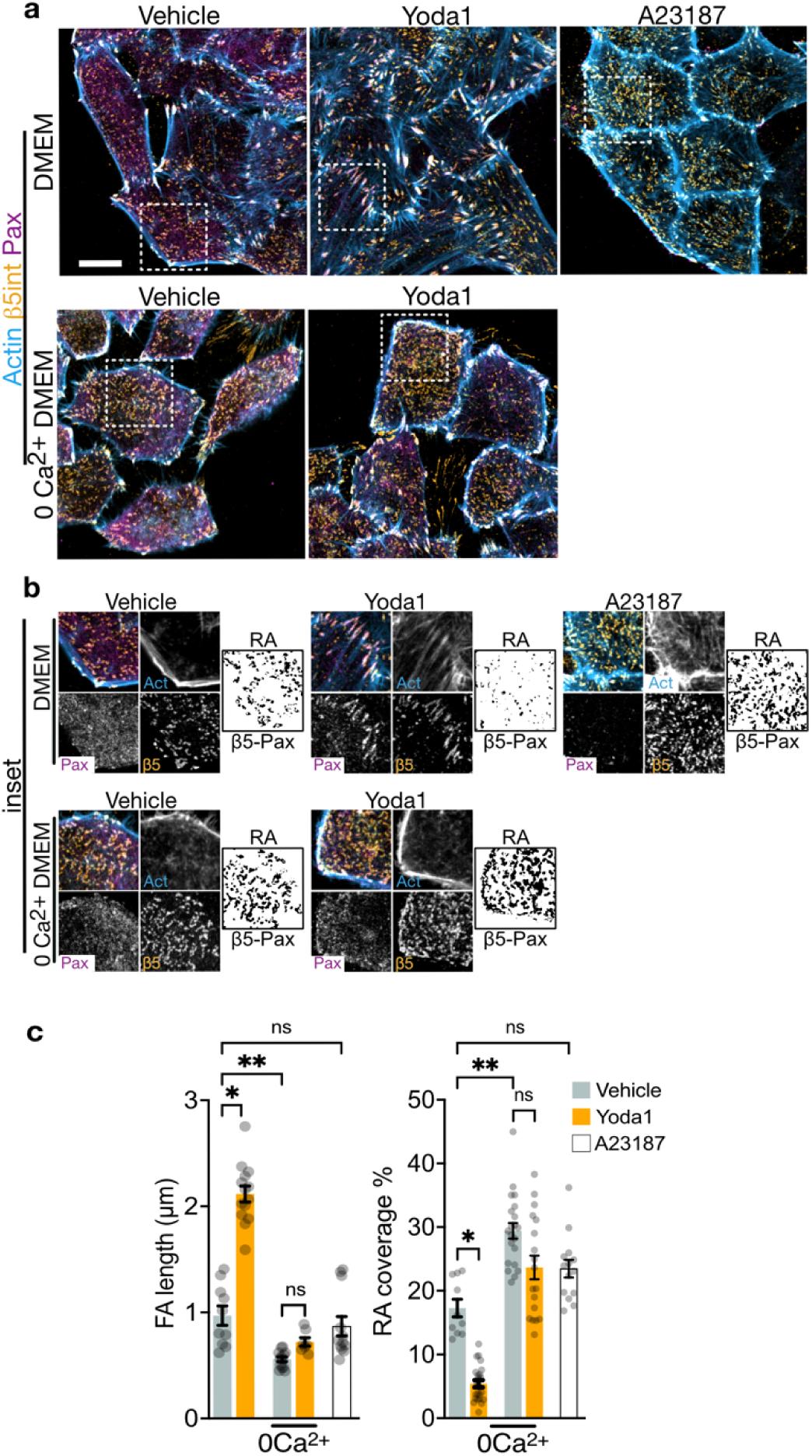
FA/RA balance by Piezo1 requires extracellular calcium. **a**. Representative confocal images of HeLa cells serum-starved overnight, treated with DMSO (vehicle control), 10 µM Yoda1, or 5 µM of the calcium ionophore A23187 in the presence (upper panel) or absence (lower panel) of extracellular calcium for 2 hours, and stained as indicated. White squares define the regions zoomed in b. Scale bar: 10μm. **b.** Zooms from a. RA image is the result of subtracting β5-Pax signals (for details, see materials and methods). **c.** Bar plots representing mean±SEM FA length from Pax stainings (left) or RA coverage expressed as the cell are percentage occupied by RA (right) in cells treated as indicated. N, FA: Vehicle=10, Yoda1=14, 0Ca^2+^=13, 0Ca^2+^+Yoda1=7, A23187=12; RA: Vehicle=10, Yoda1=24, 0Ca^2+^=21, 0Ca^2+^+Yoda1=18, A23187=14 cells from 3 experiments. Each point in graphs c represents a cell. n.s., non-significant; *p<0.05; **, p<0.01; *** p<0.001 in ANOVA followed by Kruskal-Wallis with Dunn’s correction.

### Piezo1 signals via Fyn kinase and RhoGEF VAV2 to balance FA and RA

Given that Src family tyrosine kinases (SFK) link Ca^2+^ changes to RhoA, control both FAs and RAs^17,38,40,41^, and link Piezo1 to CME^24^, we next investigated their role in Piezo1-mediated FA/RA balance. We focused on Fyn, a Ca^2+^-activated SFK that links mechanical and chemical signals to RhoA activation and FA and stress fiber formation^38,42–45^, and on VAV2, a widely expressed Rho guanine nucleotide exchange factor (RhoGEF) activated by Fyn-mediated Tyr phosphorylation that, like Piezo1, regulates CME and FA formation^38,46–49^ (Fig. 4a). Due to the lack of Fyn-specific inhibitors, we relied on siRNA-mediated Fyn knockdown. Immunoblots using an antibody against Tyr172-phosphorylated VAV2 (pVAV2), an indicator of active VAV2 in cell culture and *in vivo*^46,50^, revealed increased pVAV2 levels within 5 minutes of Yoda1 treatment, whereas Fyn knockdown reduced basal pVAV2 levels and, importantly, suppressed Yoda1-induced pVAV2 increases (Fig. 4b).

**Figure 4.**
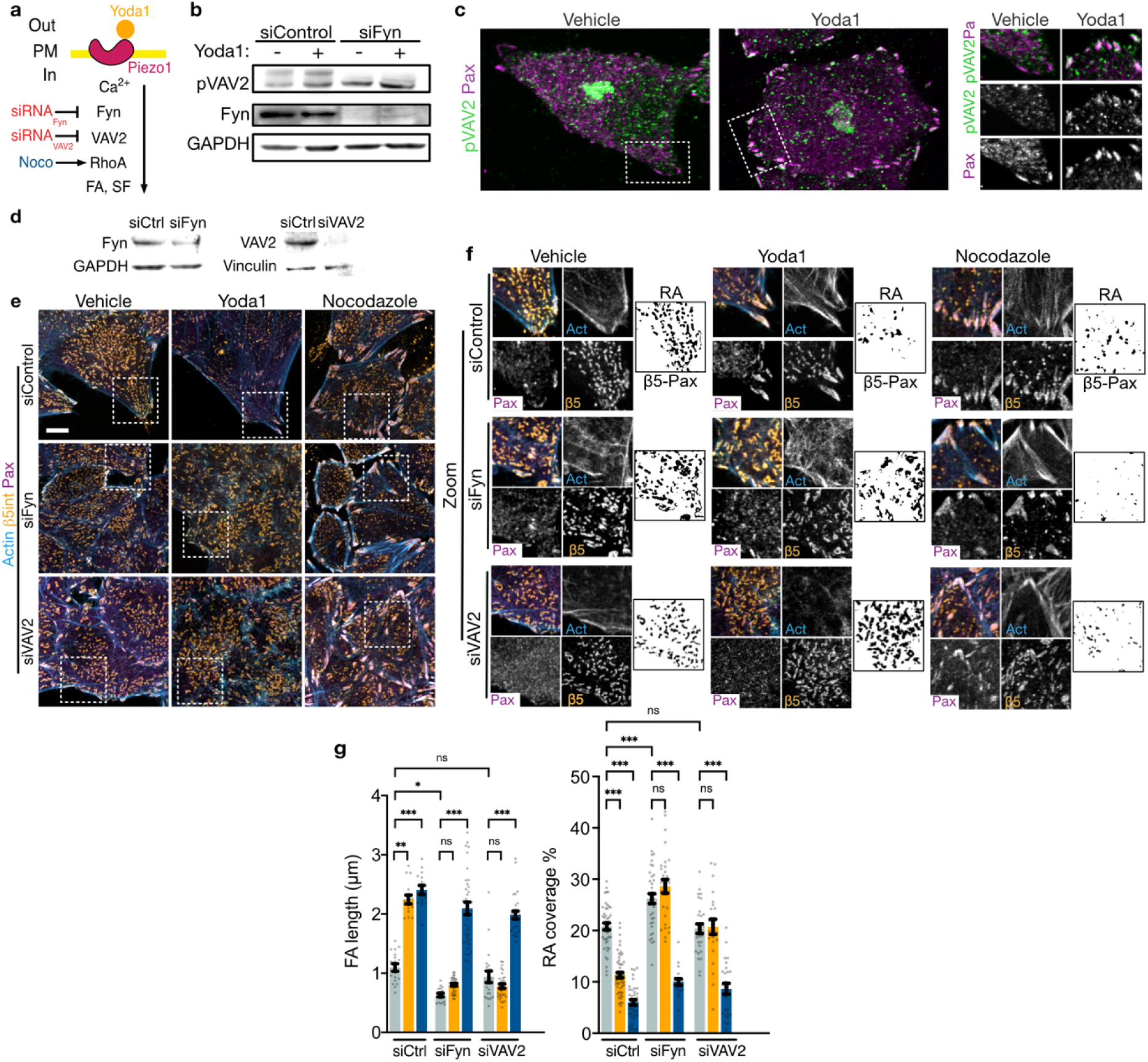
Piezo1 signals via Fyn kinase and RhoGEF VAV2 to balance FA and RA. **a.** Pathway schematic. Extracellular calcium entry via Piezo1 activates the Src family kinase Fyn. Fyn activates RhoGEF VAV2 via phosphorylation on Tyr 172. Active phosphoVAV2 exchanges GDP to GTP on RhoA, activating it. Active RhoA promotes actin polymerization and FA growth. Fyn and VAV2 were targeted with specific siRNA. RhoA can be directly activated by Nocodazole, used as a positive control. **b.** Immunoblot in lysates from cells treated with siRNA for 48h prior to overnight serum-starvation, 5-min treatments with DMSO (vehicle control) or 10μM Yoda1, and lysis. Yoda1 increases pVAV2 levels, and this is prevented by Fyn knockdown. GAPDH used as loading control. **c.** Representative confocal images of HeLa cells serum-starved overnight, treated for 2 hours with DMSO (vehicle control), 10 µM Yoda1, and stained with antibodies against Paxillin as FA marker and pVAV2. **d.** Immunoblot in lysates from cells treated with indicated siRNA for 48h prior to overnight serum-starvation and lysis. GAPDH and Vinculin used as loading controls. **e.** Representative confocal images of HeLa cells transfected with siRNA for 48h, serum-starved overnight, treated for 2 hours with DMSO (vehicle control), 10 µM Yoda1, or 25µM Nocodazole (positive control) and stained as indicated. RA image is the result of subtracting β5-Pax signals (for details, see materials and methods). White squares define the regions zoomed in f. **f.** Zooms from d. RA image is the result of subtracting β5-Pax signals (for details, see materials and methods). **g.** Bar plots representing mean±SEM FA length from Pax stainings (left) or RA coverage expressed as the cell are percentage occupied by RA (right). Each point represents a cell. N, FA: Vehicle=18 (siControl) 17 (siFyn) 21 (siVAV2), Yoda1=15 (siControl) 34 (siFyn) 31 (siVAV2), Nocodazole=15 (siControl) 39 (siFyn) 27 (siVAV2) cells, RA: Vehicle=44 (siControl) 39 (siFyn) 29 (siVAV2), Yoda1=57 (siControl) 28 (siFyn) 23 (siVAV2), Nocodazole=40 (siControl) 20 (siFyn) 26 (siVAV2) cells from 3 experiments. n.s., non-significant; *p<0.05; **, p<0.01; *** p<0.001 in ANOVA followed by Kruskal-Wallis with Dunn’s correction.

These experiments revealed that Piezo1 promotes Fyn-dependent VAV2 phospho-activation. In addition, immunostainings revealed pVAV2 accumulation at FAs in response to Yoda1 treatments (Fig. 4c), suggesting a mechanism for localized VAV2-RhoA activation, actin polymerization, and FA growth^51,52^. Having established that Piezo1 promotes VAV2-activating Tyr172 phosphorylation via Fyn and pVAV2 accumulation at FAs, we used siRNA-mediated knockdown to evaluate Fyn and VAV2 involvement in Piezo1-dependent FA/RA balance (Fig. 4d). Under basal conditions, Fyn knockdown reduced FAs and increased RA coverage, whereas VAV2 knockdown did not, suggesting that basal FA/RA balance requires Fyn but not VAV2. However, both knockdowns suppressed Yoda1-induced FA/RA changes without affecting responses to nocodazole (Fig. 4e-g). We complemented our VAV2 knockdown experiments with vilanterol, a recently described SFK-VAV2-RhoA axis inhibitor that binds to VAV2 and prevents its Tyr172 phosphorylation in cell culture and *in vivo*^50^. Vilanterol phenocopied VAV2 knockdown, as it did not alter FA length or RA area under basal conditions, and it suppressed Yoda1- but not nocodazole-induced FA growth and RA loss (Fig. S3a-c). Together, these experiments rule out nonspecific responses caused by disruption of Fyn or VAV2 and identify a Fyn-VAV2 axis controlling RhoA-dependent FA/RA balance downstream of Piezo1 activation.

### Piezo1 activates N-WASP via Fyn to disassemble RAs

Our experiments show that Piezo1 promotes RhoA-dependent FA growth, while reducing RA coverage, but do not explain how this occurs. Thus, we next investigated how Piezo1 might dissociate RAs. N-WASP promotes branched actin nucleation by Arp2/3 at the periphery of RAs, driving their disassembly into clathrin-coated endocytic vesicles^5,7,14^. However, little is known about what triggers N-WASP to dismantle RAs. Given that SFK, including Fyn, bind and activate N-WASP via Tyr 256 phosphorylation (pN-WASP)^53–55^, we tested if Piezo1 activation promotes N-WASP phosphorylation (Fig. 5a). We found that Yoda1 increased pN-WASP levels in a Fyn-dependent manner, confirming that Piezo1 promotes Fyn-dependent N-WASP phosphoactivation (Fig. 5b). Additionally, Arp2/3 inhibition with CK666 prevented Yoda1-induced, but not nocodazole-induced FA growth and RA shrinkage or basal FA/RA balance (Fig. 5c-e). Thus, Arp2/3 activity likely acts downstream of specific external signals like Piezo1 activation to drive cytoskeletal and adhesion reorganization, whereas nocodazole-induced microtubule depolymerization and RhoA activation may act in parallel. Together, our results support a model where Piezo1 signaling requires Fyn-dependent N-WASP phosphorylation and downstream Arp2/3-mediated branched actin nucleation to disassemble RAs.

**Figure 5.**
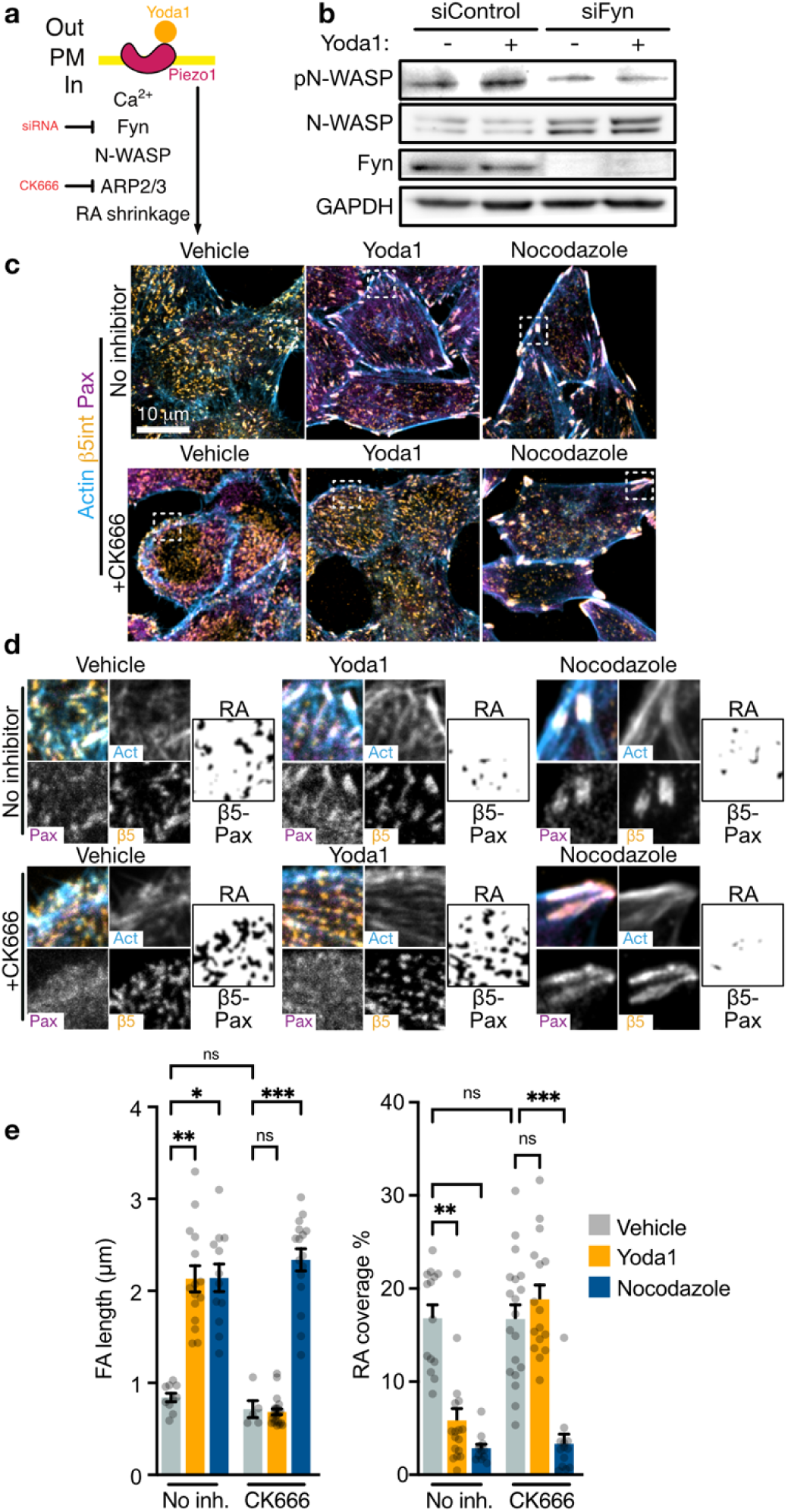
Piezo1 activates N-WASP via Fyn to disassemble RAs. **a**. Pathway schematic. Extracellular calcium entry via Piezo1 activates the Src family kinase Fyn. Fyn activates Ν-WASP via phosphorylation on Tyr 256. Active phosphoN-WASP activates Arp2/3. Active Arp2/3 promotes branched actin polymerization, providing forces for RA disassembly. Fyn was targeted with specific siRNA. CK666 was used to inhibit Arp2/3. **b.** pN-WASP immunoblot in lysates from cells treated with siRNA for 48h prior to overnight serum-starvation, 5-min treatments with DMSO (vehicle control) or 10μM Yoda1, and lysis. GAPDH used as loading control. **c.** Representative confocal images of HeLa cells stained as indicated after overnight serum starvation followed by 2h treatments with DMSO (vehicle control), 10 µM Yoda1, or 25µM Nocodazole (positive control) in the presence or absence of 100μΜ CK666 (added 30min earlier). White squares define the regions zoomed in d. Scale bar: 10μm**. d**. Zooms from c. RA image is the result of subtracting β5-Pax signals (for details, see materials and methods). **e.** Bar plots representing mean±SEM FA length from Pax stainings (left) or RA coverage expressed as the cell are percentage occupied by RA (right). Each point represents a cell. N, FA: Vehicle=10, Yoda1=15, Nocodazole=12, CK666=5, CK666+Yoda1=22, CK666+Nocodazole=16; RA: Vehicle=14, Yoda1=17, Nocodazole=12, CK666=19, CK666+Yoda1=16, CK666+Nocodazole=13 cells from 3 experiments. n.s., non-significant; *p<0.05; **, p<0.01; *** p<0.001 in ANOVA followed by Kruskal-Wallis with Dunn’s correction.

### Piezo1 enables EGFR CME mechanoadaption

RA disassembly contributes to clathrin-coated vesicle formation, and its inhibition impairs CME and signaling of specific transmembrane receptors like EGFR^9,14,17,56^. Importantly, N-WASP-Arp2/3-mediated branched actin nucleation is essential for CME under high membrane tension to enable membrane deformation required for endocytic vesicle formation. As a result, N-WASP or Arp2/3 disruption impairs CME under mechanically challenging conditions (hypotonicity, stiff substrates), likely due to excessive of RA stability^8,20–22,56,57^. However, how membrane tension activates N-WASP-Arp2/3 to license CME remains unclear. Given that membrane tension directly activates Piezo1^25,28^ and Piezo1 promotes RAs disassembly via N-WASP-Arp2/3, we hypothesized that Piezo1-dependent RA disassembly is essential for CME mechanoadaption.

To test this, we studied EGFR endocytosis in response to its extracellular soluble ligand EGF, a process known to involve CME form disassembling RAs disassembly^12,17,24,56,58^. Given that matrix stiffness regulates both Piezo1 activity and EGFR internalization^12,39^, we compared pEGFR (Tyr1173) internalization in control and Piezo1-knockdown cells grown on soft (1.5kPa hydrogel) or stiff (glass) substrates and treated with physiological EGF doses (1ng/mL) for 15 min^24,58^ (Fig. 6a). On soft substrates, pEGFR internalization was similar between control and Piezo1-knockdown cells. On glass, pEGFR internalization was reduced in both cell types, consistent with extracellular stiffness hindering CME^12,56^ (Fig. 6b-d). However, pEGFR internalization loss was greater in Piezo1-knockdown cells (Fig. 6b-d, right). Clathrin-knockdown confirmed that at this EGF dose, pEGFR internalization is CME-dependent^58^ (Fig. S4a-d). However, neither Piezo1 nor clathrin knockdown suppressed pEGFR internalization on glass at high EGF doses (100ng/mL), consistent with it being clathrin-independent^58^ (Fig. S4b-d, right). These experiments rule out general endocytic or EGFR defects in Piezo1-knockdown cells and demonstrate that Piezo1 is necessary for canonical (ligand-triggered, CME-mediated^24,58^) EGFR internalization in stiff environments, establishing a specific and essential role for Piezo1 in EGFR CME mechanoadaption^9,12,13^.

**Figure 6.**
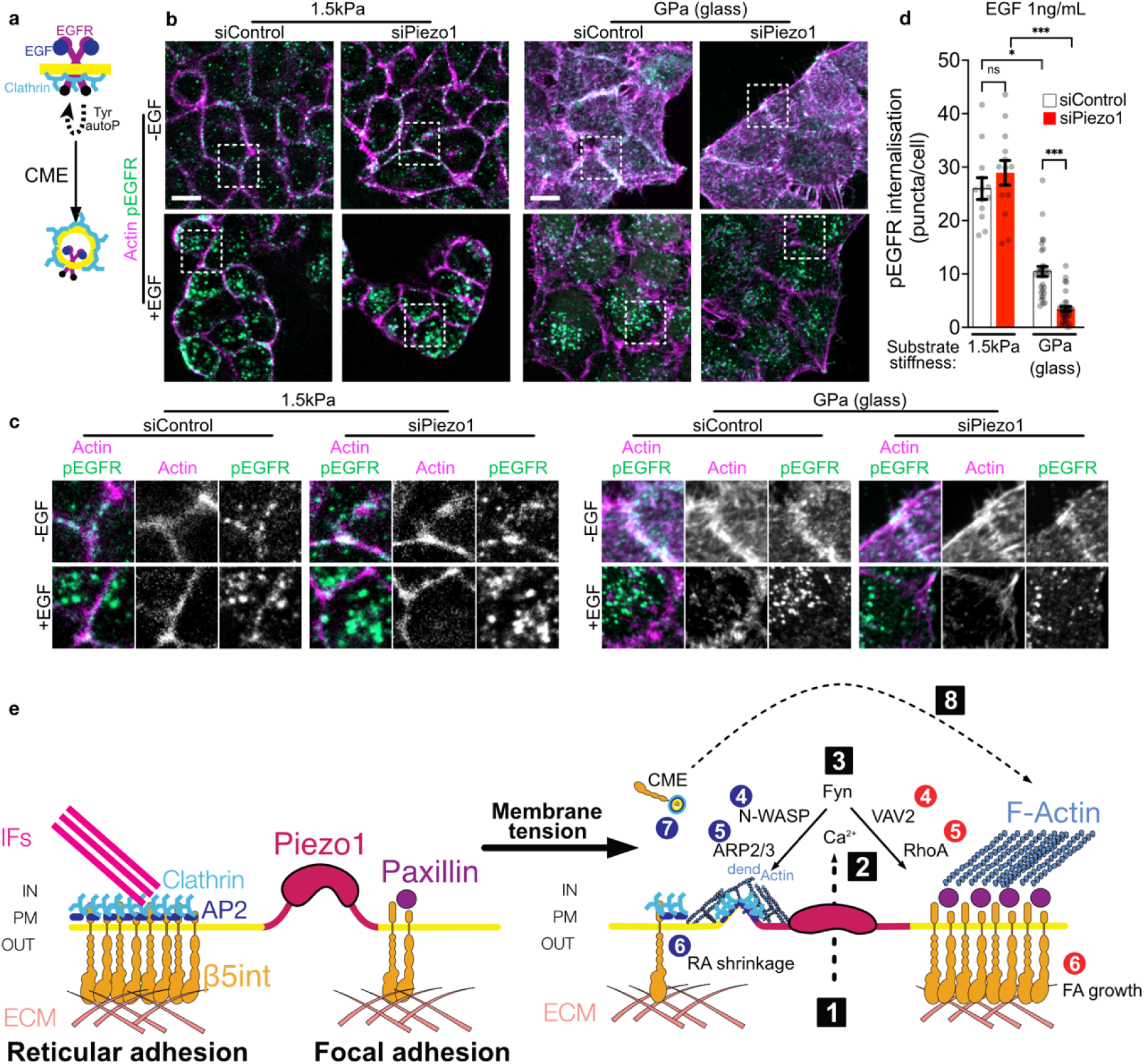
Piezo1 enables EGFR CME mechanoadaption. **a.** EGF binding promotes EGFR autophosphorylation on cytoplasmic Tyr residues, which recruits endocytic and adaptors and effectors that determine signaling output and EGFR internalization by clathrin-mediated endocytosis (CME). **b.** Representative confocal images of HeLa cells grown on vitronectin-coated hydrogels (1.5kPa stiffness) or on glass (stiffness in the GPa range), after 48h of siRNA transfection, serum-starved overnight, treated with 1ng/mL EGF for 15min, and stained as indicated. White squares define the regions zoomed in c. Scale bar: 10μm. **c.** Zooms from b. **d.** Bar plots representing mean±SEM pY1173-EGFR puncta per cell. Each point represents a cell. N, 1.5kPa: siControl=12, siPiezo1=13; Glass: siControl=31, siPiezo1=34 cells from 3 siRNA transfections. n.s., non-significant; *** p<0.001 in ANOVA followed by Kruskal-Wallis with Dunn’s correction. **e.** Model of FA/RA balance by Piezo1. Membrane tension increases activate extracellular Ca^2+^ entry via Piezo1 (1), which activates the Src kinase Fyn (2). Here, two branches split: in a first branch (blue circles), activates N-WASP via Tyr 256 phosphorylation (3). In turn, N-WASP promotes Arp2/3-mediated dendritic actin (dActin) polymerization (4 and 5), supplying energy to disassemble RA (6) and deform the plasma membrane to generate endocytic vesicles for CME (7). In the other branch (red circles), Fyn activates VAV2 via Tyr172 phosphorylation (4). In turn, VAV2 activates RhoA-dependent FA and stress fiber formation (5 and 6). In this model, RA dissolution could internalize β5 integrin via CME, enabling the integrin recruitment (8) observed during FA growth.

## Discussion

Most mechanobiology studies have focused on FAs while overlooking RAs, and while RA studies examine their role in CME, most have not addressed their interplay with FAs. Our work bridges these two fields by uncovering a previously unrecognized mechanical balance between FAs and RAs under Piezo1 control. Piezo1-dependent Ca²⁺ entry activates SFKs including Fyn, which bifurcates into two phospho-triggered actin polymerization pathways: stress fiber and FA growth via pVAV2-RhoA, and RA disassembly to enable CME via pN-WASP-Arp2/3. This bifurcation provides a mechanism by which a common mechanical stimulus can coordinate opposing FA and RA responses while supporting EGFR CME through a shared mechanosensor, Piezo1. It also reveals an unrecognized arm of Piezo1 actin control: beyond its established promotion of linear F-actin via RhoA^37,38^, we show that Piezo1 drives branched filament nucleation via N-WASP-Arp2/3, providing a molecular link between membrane tension and the actin polymerization that licenses CME under mechanical load^5,14,20,21^. Our work also expands the known functions of Piezo1 beyond mechanosignaling by demonstrating its requirement for ligand-induced EGFR internalization on stiff substrates (Fig. 6c), establishing an essential role for Piezo1 in classical receptor-mediated signaling, a largely overlooked dimension of Piezo1 studies, focused on mechanically triggered signaling.

A key question raised by our findings is how a single Piezo1-dependent Ca²⁺ signal controls competing actin networks: linear F-actin via RhoA and branched actin via Arp2/3. Our ionophore experiments show that unspecific Ca²⁺ entry fails to reproduce Yoda1-induced FA growth and RA loss, suggesting that the spatial context of Ca²⁺ influx, not its magnitude *per se*, is essential for FA/RA balance. We propose that Ca²⁺ influx via activated Piezo1 locally activates Fyn, with downstream events depend on the adhesion type: at FAs, Fyn phosphorylates VAV2 to activate RhoA, causing linear actin polymerization, whereas at RAs peripheries, Fyn phospho-activates N-WASP to drive Arp2/3-mediated branched actin nucleation. Given that β1 integrin ligation by fibronectin activates Fyn, our results may also help explain the ability of fibronectin-bound β1-integrin to promote RA disassembly^10,59^. FA/RA balance may also be imposed by adhesions sharing a limited set of structural components: β5 integrin recruitment to growing FA could drain it from RAs, contributing to their disassembly^2,29,60,61^. Simultaneously, this would release endocytic players from RAs, freeing a pool for CME. In addition, Piezo1-driven Ca²⁺ influx may activate calpain, a Ca²⁺-dependent protease involved in adhesion remodeling, which could reinforce integrin redistribution^38,62–64^. In parallel to actin-based mechanisms, two additional biochemical switches may trigger structural rearrangements within the RA lattice to promote its dissolution. First, Ca²⁺ binding to clathrin light chains induces conformational changes that promote clathrin coat curvature. Therefore, Piezo1-driven Ca²⁺ influx may directly destabilize the flat clathrin lattice of RAs^65,66^. Second, tyrosine phosphorylation of RA structural components including clathrin and β5-integrin may trigger lattice remodeling that cooperates with N-WASP-Arp2/3 to complete RA disassembly^17,32,34^. Consistent with this, phospho-tyrosine switches under Src and Activated Cdc42-Associated Kinase (ACK) control have recently been shown to promote RA disassembly and CME of specific receptors, including EGFR^17,34,67^. Together, these findings suggest that tyrosine phosphorylation of RA components is a general dissolution trigger engaged by multiple stimuli, including mechanical forces via Piezo1 and soluble ligands like EGF via EGFR.

Dividing cells disassemble FAs and adhere to the underlying matrix via RAs, whereas migrating cells disassemble RAs and form FAs that propel the cell by transmitting actomyosin forces to the ECM^4,10^. Our work shows that these changes are hardwired to be balanced via Piezo1. In dividing cells, actomyosin is a major constituent of the cell cortex, and thus it is not available for stress fiber formation. Simultaneously, Piezo1 relocates to the midbody, where it is transiently activated by mechanical forces to control timely membrane scission and cytokinesis^68^. We propose that actomyosin and Piezo1 relocation to the cortex and the midbody, respectively, cooperate to inhibit the Piezo1-driven FA/RA axis, shifting the balance towards FA shrinking and RA growth. This shift directly affects CME: the sequestration of actin in the cortex and endocytic components at RAs helps explain mitotic CME shutdown, which isolates dividing cells from signaling^69,70^. Beyond division, Piezo1 senses crowding-induced mechanical forces to trigger cell extrusion from epithelial monolayers, which requires coordinated adhesion remodeling and likely CME-dependent clearance of transmembrane receptors and junction proteins from the extruding cell membrane^71,72^. Whether the axis identified here operates in these settings will be the subject of future work.

We had previously identified essential roles for Piezo1/2 in RhoA activation by mechanical signals like confinement or matrix rigidity but had not attributed the link to any specific GEF^37,38^. Here we identify VAV2 as the GEF activating RhoA downstream of Piezo1 and Fyn. Interestingly, head and neck squamous cell carcinoma requires VAV2- and RhoA-driven YAP-dependent transcriptional programs, and constitutively active VAV2 mutants are oncogenic via RhoA-dependent actomyosin and FA rearrangements^47,73^. This underscores that cancer driver mutations are not oncogenic per se but require mechanosignaling to drive tumor growth^38,74,75^. Our work adds a new layer to this picture: beyond driving oncogenic signaling (e.g. VAV2, EGFR, YAP)^73,74,76,77^, the Piezo1-FA/RA axis controls the endocytic capacity of cancer cells and directly determines how efficiently they clear surface receptors like EGFR via CME. This is clinically relevant because CME-mediated EGFR clearance is a recognized mechanism of cancer resistance to anti-EGFR monoclonal antibodies. Tyrosine kinase inhibitors (TKI) are used as anti-CME adjuvants, but cancer cells develop additional resistance mechanisms that, unfortunately, promote relapse and patient death^78–80^. Therefore, targeting the Piezo1-RA-CME axis to reduce endocytic capacity may offer a complementary strategy to overcome TKI resistance in prevalent breast, lung, and colon cancers^81^.

EGFR is not the only receptor transiting through RAs; Hepatocyte Growth Factor Receptor (HGFR) and G protein-coupled receptors including Lysophosphatidic Acid Receptor 1 (LPAR1) and C-C Chemokine Receptor 5 (CCR5) accumulate and internalize from RAs, causing signaling outputs that differ depending on whether they signal from the plasma membrane or within the endomembrane system after internalization^14,56,82^. Therefore, Piezo1-dependent CME control could shape the signaling output of several receptors that regulate differentiation, proliferation and migration, all essential functions in development, homeostasis and disease. In addition, CCR5 and β5 integrin mediate, respectively, HIV and Zika virus entry into host cells^82–84^. Although RAs involvement in these processes remains unstudied, this potential link between Piezo1 and virus entry may also be targetable in the clinic.

Our work identifies Piezo1 as the mechanosensor that co-regulates cell-matrix adhesion and CME, integrating environmental mechanical properties with membrane and adhesion remodeling and receptor internalization^85,86^. We further show that Piezo1 loss selectively impairs ligand-triggered, CME-mediated EGFR internalization on stiff but not soft substrates, revealing a stiffness-dependent vulnerability in receptor trafficking that could lead to more precise targetable treatments.

## Materials and methods

Unless noted, reagents were obtained from Thermo Fisher Scientific.

### Cell culture

Non-verified HeLa and U2OS cells were grown (incubator, 37°C, 5% CO_2_) in DMEM (31966021) with 10% FBS (10270106) and 1% Pen/Strep (15070063). For all experiments, cells were seeded in growth medium. After 24h, growth medium was washed twice with PBS (14190250) and replaced by starvation medium (DMEM+1%Pen/Strep, without FBS) for 24h before cell treatment. Periodic PCR tests for Mycoplasma contamination (Sartorius, 20-700-20) were negative.

### siRNA transfection and culture

30-50% confluent cells were transfected with 10nM siRNA (Horizon Discovery, non-targeting control: D-001810-10-05; Piezo1: L-020870-03-0020; Fyn:_L-003140-00-0005; VAV2: L-005199-00-0005; Clathrin Heavy Chain 1: L-004001-01-0020) with Lipofectamine RNAiMax (13778) in Optimem (31985062), following manufacturer’s instructions. After 48h, cells were serum starved overnight to perform experiments 72h after transfection.

For hydrogel experiments, cells were split 24h after transfection and seeded on vitronectin-(A14700) coated 1.5 kPa hydrogels on glass bottom 35mm petri dishes (ibidi, 81291), grown for 24h and later serum-starved overnight to perform experiments 72h after transfection.

### U2OS-σAdaptin-GFP line generation

The generation of this cell line is described in detail in previous published work^10^.

### Chemicals and treatments for FA/RA balance experiments

All treatments were prepared reusing starvation medium supernatants. 24h after serum starvation, cells were treated for indicated durations with DMSO (276855), 10µM Yoda1 (Tocris, 5586), 25µM Nocodazole (Sigma, M1404), 1µM A23187 (Biovision, 1501-10), all reconstituted in DMSO, 1μg/mL Rho Activator (Cytoskeleton Inc, CN03) or 2ng/mL EGF (Peprotech, 400-25), both reconstituted in sterile deionized H_2_O. Inhibitor experiments included a 30-min pre-treatment with 10µM Vilanterol (Sigma, SML3389, reconstituted in DMSO), 1μM GsMTx4 (Tocris, 4912), or 0.5μg/mL Rho Inhibitor (Cytoskeleton Inc, CT04-A) (both reconstituted in sterile deionized H_2_O). For 0Ca^2+^ experiments, Ca^2+^-free DMEM (21068-028) was supplemented with 1% Pen/Strep and 1.2mM MgCl_2_ (Sigma, M8266) to avoid integrin inhibition.

### EGFR internalization experiments

siRNA-transfected HeLa cells grown on glass coverslips were serum-starved and treated with deionized water (dH_2_Ο) or 1ng/mL EGF (PeproTech,400-25, reconstituted in dH_2_Ο) for 15min in the cell culture incubator (37°C, 5%CO_2_).

### Immunostaining FA/RA balance and EGFR internalization experiments

After treatments, samples were rinsed twice, fixed with 4% PFA (28908, 20min, 37°C), permeabilized with 0.5% TritonX-100 (Sigma, X-100, 5min, RT), stained with and secondary antibodies, and mounted with Fluoromount-G (004958-02). Primary antibodies (2h, RT or overnight, 4°C): αvβ5 (Sigma, zrb1191, 1/300), Paxillin (BD Biosciences, 610051, 1/150), α-adaptin (Abcam, ab2730, 1/50), pVAV2 (Novus, NBP3-23269, 1/150), phosphoY1173-EGFR (R&D Systems, AF1095, 1/300). Alexa Fluor^TM^-conjugated secondary antibodies (45min, RT, 1/1000): anti-Rabbit AF488 (A11008), anti-Mouse AF549 (A11005) were incubated with DAPI (62248, 1/2000) and Phalloidin-iFluor^TM^ 647 (Abcam, ab176759, 1/1000) to stain DNA and actin, respectively. Antibody dilutions were prepared in 1% BSA (Sigma, A7906). PBS was used to prepare all solutions and washings.

### Calcium imaging

HeLa cells grown on glass coverslips were loaded with Calbryte™ 520 AM (AAT Bioquest, 20651, reconstituted in DMSO) inside the cell culture incubator (30min, 37°C, 5%CO_2,_). After two rinses, coverslips were mounted on a recording chamber (Warner Instruments, 642420) and imaged by epifluorescence at room temperature on a Nikon Eclipse Ti2 microscope using GFP-compatible filter settings, a 20x air objective and 2x2 binning. 10μΜ Yoda1 was freshly prepared before each replicate from 10mM stocks. All solutions were prepared in Fluobrite DMEM (A1896701) supplemented with 20mM HEPES (15630080). Timecourse graphs depict the background-substracted fluorescence signal at each timepoint, corrected by the signal at the start of the experiment.

### Confocal imaging

Unidirectional line-scanning confocal imaging was done at room temperature on a Leica SP8 microscope with a 63x objective (ref. 11506349, numerical aperture 1.4, Plan Apo, working distance 0.14mm) and immersion oil (refractive index 1.51), with 633, 561, 488, and 405 nm lasers and two conventional photomultipliers (PMT). All settings were controlled with Leica LAS X software. PMT gain and offset were adjusted avoiding saturation and low signal cutoff and left unaltered when comparing the same staining between conditions. Frame average was 4 and resolution was 1024 x 1024 pixels. Pinhole size was 1 AU. Pixel size was 90.19x90.19nm and z-stack slice thickness was 0.3μm. Initial and final Z were defined using FA-like Paxillin signal. These settings were constant regardless of staining, treatment, or session. All image analysis was done with FIJI^87^.

#### Focal adhesion analysis

FA length was manually measured on bright Pax stainings at cell edges, as previously described^2^.

#### Reticular adhesion analysis

Thresholded and dilated Paxillin images were subtracted from thresholded αvβ5 images, resulting in a RA mask of αvβ5^+^, Pax^-^ spots. RA coverage was then calculated as (integrated density/255) and expressed as the percentage of cell area occupied by RA, as previously described^10^.

#### EGFR internalization analysis

pY1173-EGFR intracellular puncta were semi-automatically counted using a custom-built Cell Profiler^88^ pipeline shared in our previous work on Piezo1 and EGFR CME^77^. In brief, individual HeLa cells were segmented from maximum intensity projections based on DNA and actin stainings. pY1173-EGFR speckles per cell were enhanced with a feature size of 10 and were automatically counted. Data was exported into comma-separated values (CSV) for statistical analysis.

### TIRF imaging

U2OS-σAdaptin-GFP cells were imaged at 37°C in serum starvation medium supplemented 25μM HEPES. The system was a Hamamatsu sCMOS Orca flash 4 V3 camera mounted on an ONI nanoimager microscope with 647, 561, 488 and 405 lasers and an Olympus 1.49NA 100x super achromatic objective. Exposure time was 300 ms and one frame was acquired every second for up to 30min.

### Cell unroofing, platinum-replica sample processing and EM of platinum replicas

Unroofing was performed by sonication as previously described^89^. Coverslips where quickly rinsed three times in Ringer+Ca^2+^ (155 mM NaCl, 3 mM KCl, 3 mM NaH_2_PO_4_, 5 mM HEPES, 10 mM glucose, 2 mM CaCl_2_, 1 mM MgCl_2_, pH 7.2), then immersed 10 s in Ringer-Ca^2+^ (155 mM NaCl, 3 mM KCl, 3 mM NaH_2_PO_4_, 5 mM HEPES, 10 mM glucose, 3 mM EGTA, 5 mM MgCl_2_, pH 7.2) containing 0.5 mg/mL poly-L-lysine) and quickly rinsed in Ringer-Ca^2+^. Cells were unroofed by scanning the coverslip with rapid (2-5 s) sonicator pulses at the lowest deliverable power in KHMgE buffer (70 mM KCl, 30 mM HEPES, 5 mM MgCl_2_, 3 mM EGTA, pH 7.2). Unroofed cells were immediately fixed using fixative in KHMgE with 2 % PFA and 2 % glutaraldehyde for 10 to 20 min for PREM morphology. Cells were further sequentially treated with 0.5 % OsO_4_, 1 % tannic acid and 1 % uranyl acetate prior to graded ethanol dehydration and hexamethyldisilazane substitution (HMDS, Sigma). Dried samples were then rotary-shadowed with 2 nm of platinum and 5-8 nm of carbon using an ACE600 high vacuum metal coater (Leica Microsystems). The resultant platinum replicas were floated off the glass with hydrofluoric acid (5 %), washed several times on distilled water, and picked up on 200 mesh formvar/carbon-coated EM grids. Replicas on EM grids were mounted in a eucentric side-entry goniometer stage of a transmission electron microscope operated at 80 kV (model CM120; Philips) and images were recorded with a Morada digital camera (Olympus). Images were processed in Photoshop (Adobe) to adjust brightness and contrast and presented in inverted contrast. Tomograms were made by collecting images at the tilt angles up to ± 25° relative to the plane of the sample with 5° increments. Images were aligned by layering them on top of each other in Photoshop. Measurements of cup length and width were performed on high magnification PREM views using ImageJ.

### Western blot

Cells were seeded in 6-well plates and treated when 60-80% confluent. After treatment, samples were placed on ice, rinsed twice with ice-cold PBS and incubated in 100 μL of RIPA buffer (89901) supplemented with protease (78430) and phosphatase inhibitor (Merck, 524625) cocktails and EDTA for 1min on ice. Then cells were scrapped and the lysate placed in pre-labelled cold tubes in ice for 30min, vortexing every 5min. Lysates were then centrifuged (13000 rpm, 10min, 4°C) and the supernatant transferred to new tubes for protein quantification with a BCA kit (23225) and storage at -20°C.

35µg grams of protein adjusted to 10 µL in RIPA and dying (B0007) buffers were denatured at 95°C for 10 min, spun down, supplemented with sample reducing buffer (B0004), and loaded in Bis-Tris gels (4-12%, NW04120BOX). A well with 5 µL of a pre-stained protein standard (LC5925) was used to track protein separation during electrophoresis at 110V for 2h in MOPS SDS buffer (B0001). Proteins were then transferred to nitrocellulose membranes (IB23002) with a dry transfer device (IB21001) at 25V for 8min. Membranes were blocked for 1h at RT with 5% BSA (for phosphorylated targets) or with 5% non-fat powder milk. Primary antibodies were incubated overnight at 4°C with constant rotation, followed by three rinses and secondary antibody incubation for 1h at RT, 2 additional rinses and 1-min incubation with ECL substrate (32209) and imaging. Analyze/Gels tool in Fiji was used for band density quantification. 0.1% Tween-20 (Sigma, P137) TBS (TTBS) was used for blocking solutions and washings. Antibodies against phosphorylated targets (pVAV2, Novus Biological, NBP3-23269; pN-WASP, PA5-143792, Fyn, BD Biosciences 610163) were prepared 1/1000 in 5% BSA-TTBS. The rest (VAV2, Cell Signaling, 2848T; N-WASP, Abcam, ab126626; GAPDH, abcam, ab8245; Vinculin, Proteintech, 26520-1-AP) were prepared in 5% non-fat dry milk-TTBS at 1/1000 dilutions, except GAPDH, prepared at 1/2000. HRP-conjugated anti-Rabbit (65-6120) or anti-Mouse (Cell Signaling, 7076) secondary antibodies were prepared 1/2000 in 5% non-fat dry milk-TTBS.

### Statistical analysis

All graphs and statistical analyses were done with Prism 11.0.2 (GraphPad Software). When comparing more than two conditions, one-way ANOVA was followed by Kluskal-Wallis nonparametric post-hoc test with Dunn’s correction for multiple comparisons. If data distribution were normal, one-way ANOVA was followed by Mann-Whitney. For 1-to-1 comparisons, significance was assessed with Welch’s test. The threshold for statistical significance was two-tailed p<0.05. N and significance level values are reported in figure legends.

## Data availability

Raw data are available upon request to C.P.-P.

## Acknowledgements

We thank all the members of the Laboratory of Molecular Physiology (UPF) and the Rosenblatt group (KCL and The Francis Crick Institute), F. Brodsky, M. Redd, A. Elosegui-Artola, and M. Rigau for helpful discussions and encouragement. We thank C. Plata-Fernández for excellent technical support and R. Garcia Olivas at the PRBB’s Advanced Light Microscopy Unit for assistance with confocal microscopes.

This work was supported by a long-term fellowship (LT000654/2019-L) from the Human Frontier Science Program organization and a Marie Skłodowska-Curie Fellowship (898067) from the European Union’s Horizon 2020 research and innovation program (C.P.-P.), Agence Nationale de la Recherche grant ANR-21-CE13-0018-01 (S.V.), and a Wellcome Trust Investigator Award (221908/Z/20/Z), a Cancer Research UK grant (DRCNPG-May21\100007), and an Academy of Medical Sciences Professorship (APR2\1007) (J.R.). This work was also supported by the Spanish Ministry of Science and Innovation and Agencia Estatal de Investigación plus FEDER Funds through grants PID2023-149767OB-I00 and ‘Unidad de Excelencia María de Maeztu’ CEX2024-001431-M funded by MICIU/AEI/10.13039/501100011033 and by ‘ERDF A way of making Europe’. L. A.-S. is supported by the Research Council of Finland and HiLIFE. L.H. is supported by The Finnish Cultural Foundation and the Finnish Cancer Institute.

C.P.-P. wishes to dedicate this work to Joana and Anton Pardo Rigau and in loving memory to Pepa Tomás Valiente and Toni Adam Donat.

## Author contributions

Conceptualization: C.P.-P.; Data curation: C.P.-P., O.B., S.B.M., S.V.; Formal analysis: C.P.-P., O.B., S.B.M., S.V.; Funding acquisition: C.P.-P., F.J.M., J.R., M.A.V., S.V.; Investigation: C.P.-P., D.C.B., O.B., O.L., R.M., S.B.M., V.A.; Methodology: C.P.-P., D.C.B., R.M., O.B., S.B.M., S.V.; Project administration: C.P.-P.; Resources: A.-S.L, C.P.-P., F.J.M., J.R., L.A.-S., L.H., M.A.V.; Software: C.P.-P., O.B., R.M.; Supervision: C.P.-P.; Validation: C.P.-P., D.C.B., O.B., R.M., S.B.M.; Visualization: C.P.-P.; Writing-original draft: C.P.-P.; Writing-review and editing: all authors

## Supplementary figures

**Figure S1.**
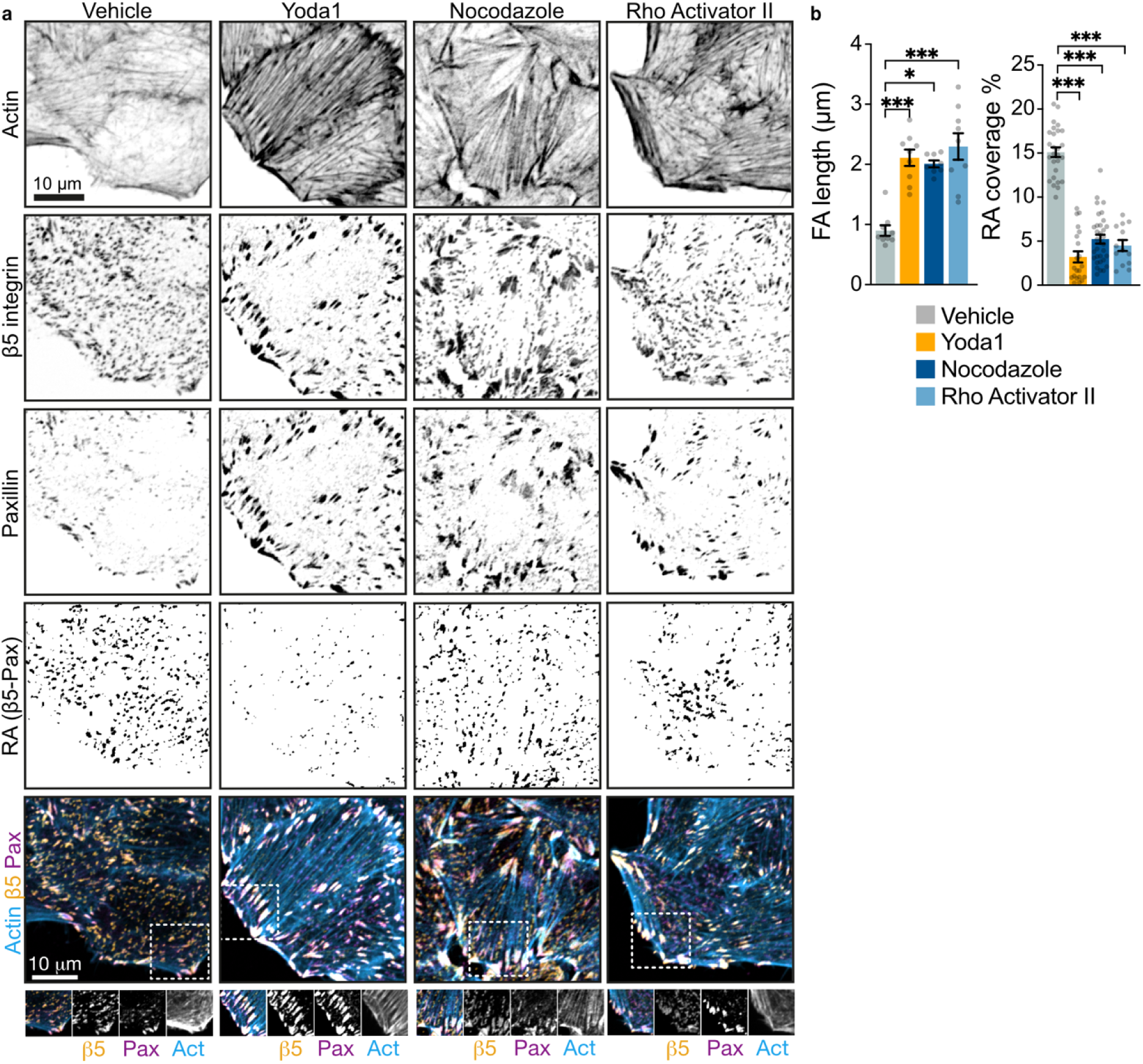
**Piezo1 balances FAs and RAs via RhoA**. **a.** Representative confocal images of HeLa cells stained as indicated after overnight serum starvation followed by 2h treatments with DMSO (vehicle control), 10 µM Yoda1, and positive controls 25µM Nocodazole and 1μg/mL RhoActivator II. RA image is the result of subtracting β5-Pax signals (for details, see materials and methods). GsMTx4 (e) was added 30min before Yoda1. Scale bar: 10μm. **b.** Bar plots representing mean±SEM FA length from Pax stainings (left) or RA coverage expressed as the cell are percentage occupied by RA (right). N, FA: Vehicle= 9, Yoda1= 9, Rho Activator= 9, Nocodazole= 8; RA: Vehicle= 27, Yoda1= 21, Rho Activator= 13, Nocodazole= 32 cells from 2 experiments. n.s., non-significant; *p<0.05; *** p<0.001 in ANOVA followed by Kruskal-Wallis with Dunn’s correction.

**Figure S2.**
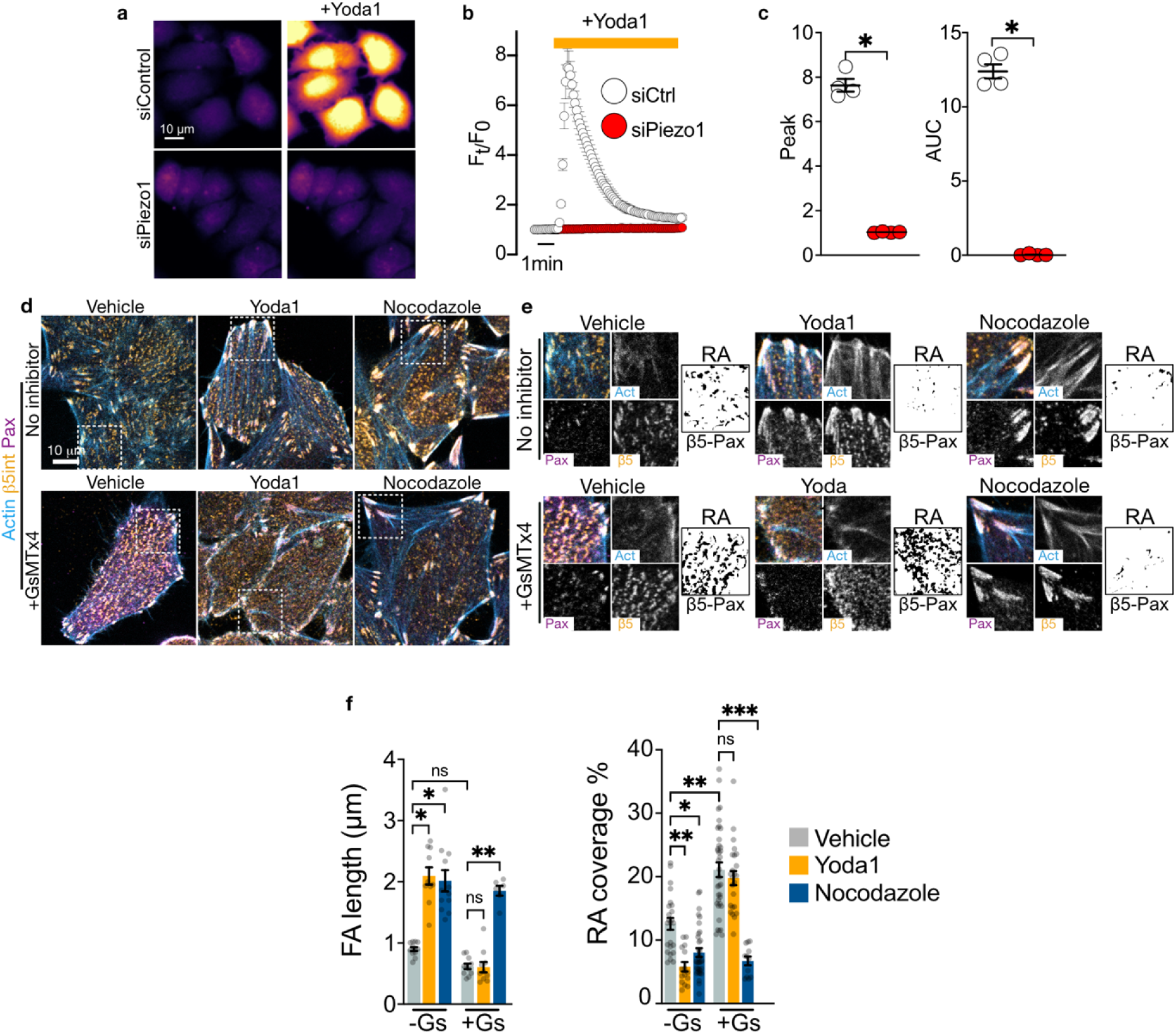
Piezo1 knockdown promotes RAs. **a.** Representative pseudo-colored stills of timelapse calcium imaging of siRNA-transfected HeLa cells 1min before and 1 min after Yoda1 treatment. Cells were transfected with siRNA for 48h and serum-starved overnight prior to imaging. **b.** Average intracellular calcium signals of siRNA-transfected HeLa cells **c.** Bar plots representing mean±SEM peak (left) and area under the curve (AUC, right) of the responses to Yoda1 (bottom). N=4 for each siRNA, from 2 independent transfections. At least 20 cells were analyzed in each experiment. **d.** Representative confocal images of HeLa cells stained as indicated after overnight serum starvation followed by 2h treatments with DMSO (vehicle control), 10 µM Yoda1, and 25µM Nocodazole (positive control) in absence or presence of 1μM GsMTx4 (added 30min earlier). White squares define the regions zoomed in h. Scale bar: 10μm. **e.** Zooms from d. **f.** Bar plots representing mean±SEM FA length from Pax stainings (left) or RA coverage expressed as the cell are percentage occupied by RA (right) in cells treated as indicated. N, FA: Vehicle=12, Yoda1=10, Nocodazole=12, GsMTx4=11, GsMTx4+Yoda1=11, GsMTx4+Nocodazole=6; RA: Vehicle=24, Yoda1=15, Nocodazole=34, GsMTx4=36, GsMTx4+Yoda1=24, GsMTx4+Nocodazole=12 cells from 3 experiments; n.s., non-significant; *p<0.05; **, p<0.01; *** p<0.001 in ANOVA followed by Kruskal-Wallis with Dunn’s correction (f) or Mann-Whitney (c) tests.

**Figure S3.**
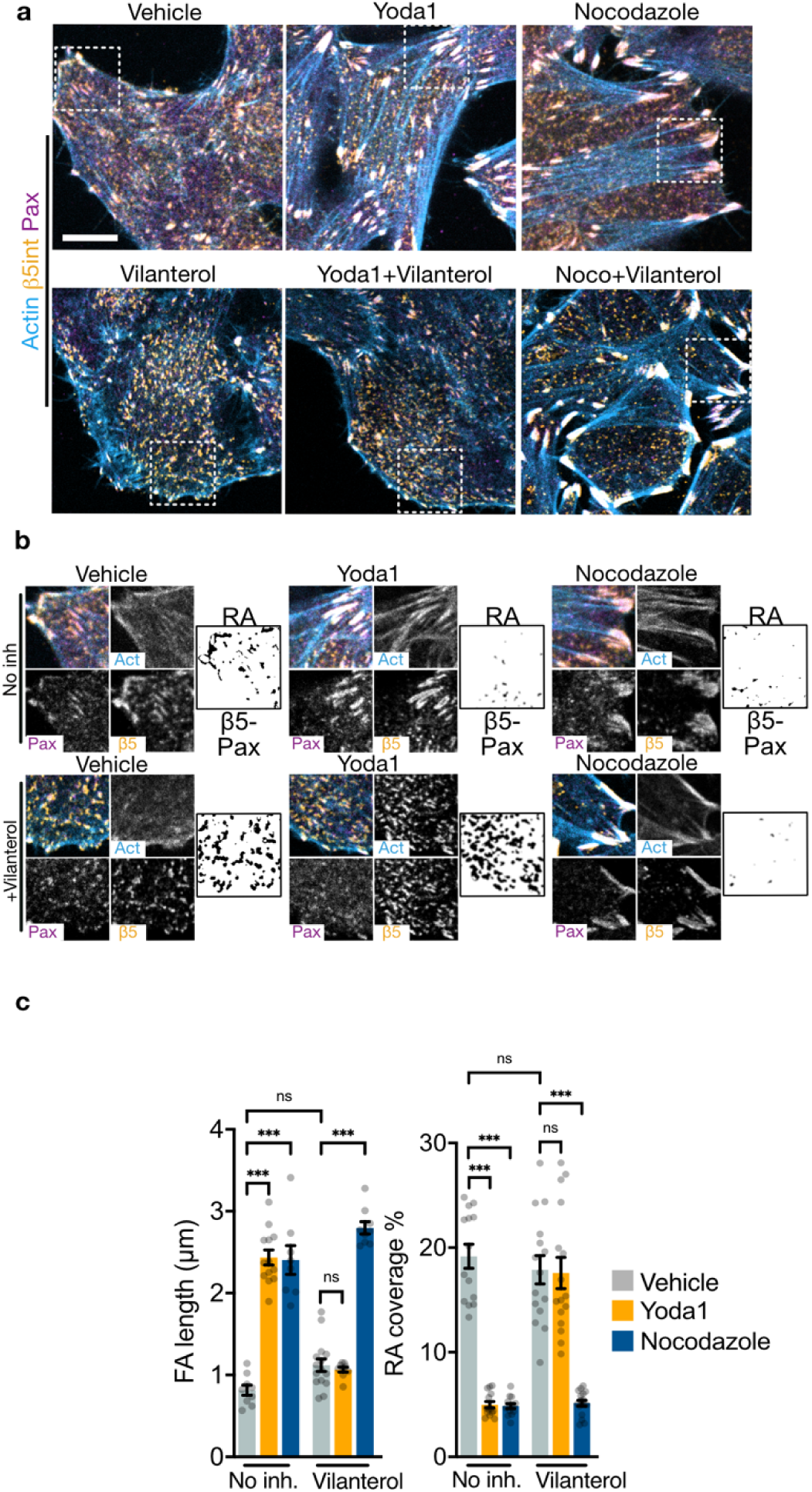
VAV2 inhibitor Vilanterol impairs FA/RA balance by Piezo1. **a.** Representative confocal images of HeLa cells stained as indicated after overnight serum starvation followed by 2h treatments with DMSO (vehicle control), 10 µM Yoda1, or 25µM Nocodazole (positive control) in the presence or absence of 10μΜ Vilanterol (added 30min earlier). White squares define the regions zoomed in b. Scale bar: 10μm**. b**. Zooms from a. RA image is the result of subtracting β5-Pax signals (for details, see materials and methods). **c.** Bar plots representing mean±SEM FA length from Pax stainings (left) or RA coverage expressed as the cell are percentage occupied by RA (right). Each point represents a cell. N, FA: Vehicle=9, Yoda1=13, Nocodazole=8, Vilanterol=15, Vilanterol+Yoda1=9, Vilanterol+Nocodazole=9; RA: Vehicle=14, Yoda1=14, Nocodazole=14, Vilanterol=15, Vilanterol+Yoda1=16, Vilanterol+Nocodazole=17 cells from 3 experiments. n.s., non-significant; *p<0.05; **, p<0.01; *** p<0.001 in ANOVA followed by Kruskal-Wallis with Dunn’s correction.

**Figure S4.**
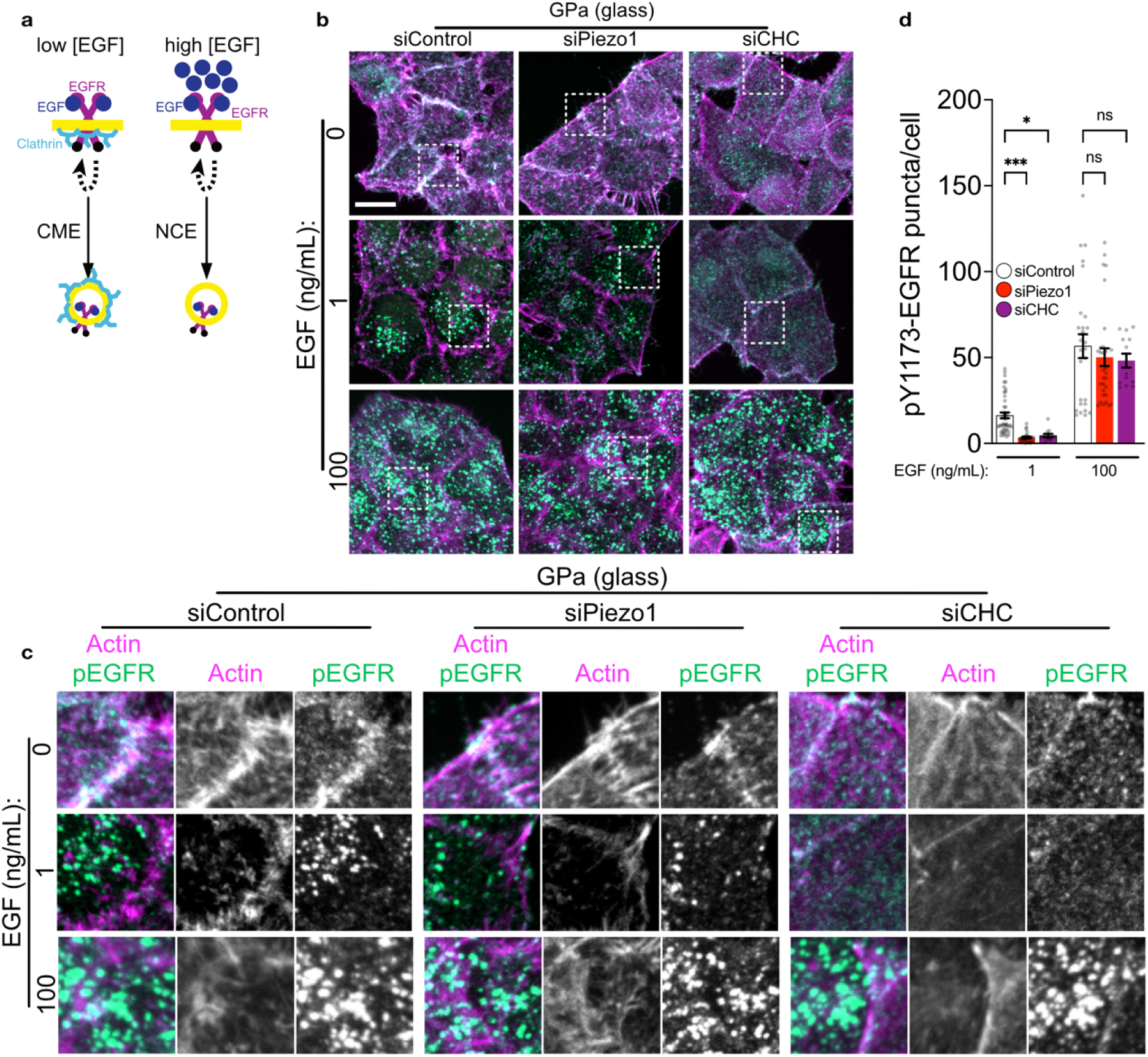
Piezo1 is required for EGFR internalization specifically by CME. **a.** Low EGF doses trigger EGFR internalization by CME, whereas high EGF doses trigger nonclathrin endocytosis (NCE)^58^. **b.** Representative confocal images of HeLa cells grown on glass (stiffness in the GPa range), after 48h of siRNA transfection, serum-starved overnight, treated with 1 or 100ng/mL EGF for 15min, and stained as indicated. White squares define the regions zoomed in c. Scale bar: 10μm. **c.** Zooms from b. **d.** Bar plots representing mean±SEM pY1173-EGFR puncta per cell. Each point represents a cell. N: siControl=45, siPiezo1=29, siCHC=12 (EGF 1ng/mL); siControl=27, siPiezo1=30, siCHC=13 (EGF 100ng/mL) cells from 3 siRNA transfections. n.s., non-significant; *p<0.05; *** p<0.001 in ANOVA followed by Kruskal-Wallis with Dunn’s correction.

## References

1. Kanchanawong, P. & Calderwood, D. A. Organization, dynamics and mechanoregulation of integrin-mediated cell–ECM adhesions. Nat Rev Mol Cell Biol 24, 142–161 (2023).

2. Elosegui-Artola, A. et al. Mechanical regulation of a molecular clutch defines force transmission and transduction in response to matrix rigidity. Nat Cell Biol 18, 540–548 (2016).

3. Ridley, A. J. & Hall, A. The small GTP-binding protein rho regulates the assembly of focal adhesions and actin stress fibers in response to growth factors. Cell 70, 389–399 (1992).

4. Lock, J. G. et al. Reticular adhesions are a distinct class of cell-matrix adhesions that mediate attachment during mitosis. Nat Cell Biol 20, 1290–1302 (2018).

5. Yang, C. et al. Actin polymerization promotes invagination of flat clathrin-coated lattices in mammalian cells by pushing at lattice edges. Nat Commun 13, 6127 (2022).

6. Vassilopoulos, S. Unconventional roles for membrane traffic proteins in response to muscle membrane stress. Current Opinion in Cell Biology 65, 42–49 (2020).

7. Franck, A. et al. Clathrin plaques and associated actin anchor intermediate filaments in skeletal muscle. MBoC 30, 579–590 (2019).

8. Saffarian, S., Cocucci, E. & Kirchhausen, T. Distinct Dynamics of Endocytic Clathrin-Coated Pits and Coated Plaques. PLoS Biol 7, e1000191 (2009).

9. Lampe, M., Vassilopoulos, S. & Merrifield, C. Clathrin coated pits, plaques and adhesion. Journal of Structural Biology 196, 48–56 (2016).

10. Hakanpää, L. et al. Reticular adhesions are assembled at flat clathrin lattices and opposed by active integrin α5β1. Journal of Cell Biology 222, e202303107 (2023).

11. Franck, A., et al. Mechanosensitive Clathrin Platforms Anchor Desmin Intermediate Filaments in Skeletal Muscle. http://biorxiv.org/lookup/doi/10.1101/321885 (2018) doi:10.1101/321885.

12. Baschieri, F. et al. Frustrated endocytosis controls contractility-independent mechanotransduction at clathrin-coated structures. Nat Commun 9, 3825 (2018).

13. Kaksonen, M. & Roux, A. Mechanisms of clathrin-mediated endocytosis. Nat Rev Mol Cell Biol 19, 313–326 (2018).

14. Leyton-Puig, D. et al. Flat clathrin lattices are dynamic actin-controlled hubs for clathrin-mediated endocytosis and signalling of specific receptors. Nat Commun 8, 16068 (2017).

15. Heuser, J. Three-dimensional visualization of coated vesicle formation in fibroblasts. The Journal of cell biology 84, 560–583 (1980).

16. Maupin, P. & Pollard, T. D. Improved preservation and staining of HeLa cell actin filaments, clathrin-coated membranes, and other cytoplasmic structures by tannic acid-glutaraldehyde-saponin fixation. Journal of Cell Biology 96, 51–62 (1983).

17. Alfonzo-Méndez, M. A., Sochacki, K. A., Strub, M.-P. & Taraska, J. W. Dual clathrin and integrin signaling systems regulate growth factor receptor activation. Nat Commun 13, 905 (2022).

18. Sigismund, S., Lanzetti, L., Scita, G. & Di Fiore, P. P. Endocytosis in the context-dependent regulation of individual and collective cell properties. Nat Rev Mol Cell Biol 22, 625–643 (2021).

19. Gauthier, N. C., Masters, T. A. & Sheetz, M. P. Mechanical feedback between membrane tension and dynamics. Trends in Cell Biology 22, 527–535 (2012).

20. Benesch, S. et al. N-WASP deficiency impairs EGF internalization and actin assembly at clathrin-coated pits. Journal of Cell Science 118, 3103–3115 (2005).

21. Boulant, S., Kural, C., Zeeh, J.-C., Ubelmann, F. & Kirchhausen, T. Actin dynamics counteract membrane tension during clathrin-mediated endocytosis. Nat Cell Biol 13, 1124–1131 (2011).

22. Merrifield, C. J., Qualmann, B., Kessels, M. M. & Almers, W. Neural Wiskott Aldrich Syndrome Protein (N-WASP) and the Arp2/3 complex are recruited to sites of clathrin-mediated endocytosis in cultured fibroblasts. European Journal of Cell Biology 83, 13–18 (2004).

23. Lappalainen, P., Kotila, T., Jégou, A. & Romet-Lemonne, G. Biochemical and mechanical regulation of actin dynamics. Nat Rev Mol Cell Biol 23, 836–852 (2022).

24. Pardo-Pastor, C. & Rosenblatt, J. Piezo1 activates non-canonical EGFR endocytosis and signaling. Sci. Adv. 9, (2023).

25. Cox, C. D. et al. Removal of the mechanoprotective influence of the cytoskeleton reveals PIEZO1 is gated by bilayer tension. Nat Commun 7, 10366 (2016).

26. Xiao, B. Mechanisms of mechanotransduction and physiological roles of PIEZO channels. Nat Rev Mol Cell Biol 25, 886–903 (2024).

27. Coste, B. et al. Piezo1 and Piezo2 Are Essential Components of Distinct Mechanically Activated Cation Channels. Science 330, 55–60 (2010).

28. Syeda, R. et al. Piezo1 Channels Are Inherently Mechanosensitive. Cell Reports 17, 1739–1746 (2016).

29. Yao, M. et al. Force- and cell state–dependent recruitment of Piezo1 drives focal adhesion dynamics and calcium entry. Sci. Adv. 8, eabo1461 (2022).

30. Syeda, R. et al. Chemical activation of the mechanotransduction channel Piezo1. eLife 4, e07369 (2015).

31. Bershadsky, A., Chausovsky, A., Becker, E., Lyubimova, A. & Geiger, B. Involvement of microtubules in the control of adhesion-dependent signal transduction. Current Biology 6, 1279–1289 (1996).

32. Zuidema, A. et al. Molecular determinants of αVβ5 localization in flat clathrin lattices – role of αVβ5 in cell adhesion and proliferation. Journal of Cell Science 135, jcs259465 (2022).

33. Bae, C., Sachs, F. & Gottlieb, P. A. The mechanosensitive ion channel Piezo1 is inhibited by the peptide GsMTx4. Biochemistry 50, 6295–6300 (2011).

34. Maib, H., Ferreira, F., Vassilopoulos, S. & Smythe, E. Cargo regulates clathrin-coated pit invagination via clathrin light chain phosphorylation. J Cell Biol 217, 4253–4266 (2018).

35. Heuser, J. The Production of ‘Cell Cortices’ for Light and Electron Microscopy. Traffic 1, 545–552 (2000).

36. Dambournet, D. et al. Genome-edited human stem cells expressing fluorescently labeled endocytic markers allow quantitative analysis of clathrin-mediated endocytosis during differentiation. J Cell Biol 217, 3301–3311 (2018).

37. Hung, W.-C. et al. Confinement Sensing and Signal Optimization via Piezo1/PKA and Myosin II Pathways. Cell Reports 15, 1430–1441 (2016).

38. Pardo-Pastor, C. et al. Piezo2 channel regulates RhoA and actin cytoskeleton to promote cell mechanobiological responses. Proceedings of the National Academy of Sciences 115, 1925–1930 (2018).

39. Pathak, M. M. et al. Stretch-activated ion channel Piezo1 directs lineage choice in human neural stem cells. Proc. Natl. Acad. Sci. U.S.A. 111, 16148–16153 (2014).

40. Eliceiri, B. P. et al. Src-mediated coupling of focal adhesion kinase to integrin **α** v **β** 5 in vascular endothelial growth factor signaling. The Journal of Cell Biology 157, 149–160 (2002).

41. Nakao, F. et al. Involvement of Src Family Protein Tyrosine Kinases in Ca^2+^ Sensitization of Coronary Artery Contraction Mediated by a Sphingosylphosphorylcholine-Rho-Kinase Pathway. Circulation Research 91, 953–960 (2002).

42. Canobbio, I. et al. The focal adhesion kinase Pyk2 links Ca2+ signalling to Src family kinase activation and protein tyrosine phosphorylation in thrombin-stimulated platelets. Biochemical Journal 469, 199–210 (2015).

43. Huang, R. Y.-J., Wang, S.-M., Hsieh, C.-Y. & Wu, J.-C. Lysophosphatidic acid induces ovarian cancer cell dispersal by activating Fyn kinase associated with p120-catenin. International Journal of Cancer 123, 801–809 (2008).

44. Thompson, W. R. et al. Mechanically activated Fyn utilizes mTORC2 to regulate RhoA and adipogenesis in mesenchymal stem cells. Stem Cells 31, 2528–2537 (2013).

45. Xu, D. et al. Involvement of Fyn tyrosine kinase in actin stress fiber formation in fibroblasts. FEBS Letters 581, 5227–5233 (2007).

46. Peng, F. et al. Mechanical stretch-induced RhoA activation is mediated by the RhoGEF Vav2 in mesangial cells. Cellular Signalling 22, 34–40 (2010).

47. Schuebel, K. E. Phosphorylation-dependent and constitutive activation of Rho proteins by wild-type and oncogenic Vav-2. The EMBO Journal 17, 6608–6621 (1998).

48. Wu, X., Suetsugu, S., Cooper, L. A., Takenawa, T. & Guan, J.-L. Focal Adhesion Kinase Regulation of N-WASP Subcellular Localization and Function. Journal of Biological Chemistry 279, 9565–9576 (2004).

49. Cowan, C. W. et al. Vav Family GEFs Link Activated Ephs to Endocytosis and Axon Guidance. Neuron 46, 205–217 (2005).

50. Zhou, T. et al. CD64^+^fibroblast-targeted vilanterol and a STING agonist augment CLDN18.2 BiTEs efficacy against pancreatic cancer by reducing desmoplasia and enriching stem-like CD8^+^T cells. Gut 73, 1984–1998 (2024).

51. Hotulainen, P. & Lappalainen, P. Stress fibers are generated by two distinct actin assembly mechanisms in motile cells. Journal of Cell Biology 173, 383–394 (2006).

52. Oakes, P. W., Beckham, Y., Stricker, J. & Gardel, M. L. Tension is required but not sufficient for focal adhesion maturation without a stress fiber template. Journal of Cell Biology 196, 363–374 (2012).

53. Suetsugu, S. et al. Sustained Activation of N-WASP through Phosphorylation Is Essential for Neurite Extension. Developmental Cell 3, 645–658 (2002).

54. Torres, E. & Rosen, M. K. Protein-tyrosine Kinase and GTPase Signals Cooperate to Phosphorylate and Activate Wiskott-Aldrich Syndrome Protein (WASP)/Neuronal WASP. Journal of Biological Chemistry 281, 3513–3520 (2006).

55. Banin, S. et al. Wiskott–Aldrich syndrome protein (WASp) is a binding partner for c-Src family protein-tyrosine kinases. Current Biology 6, 981–988 (1996).

56. Baschieri, F., Le Devedec, D., Tettarasar, S., Elkhatib, N. & Montagnac, G. Endocytosis frustration potentiates compression-induced receptor signalling. Journal of Cell Science jcs.239681 (2020) doi:10.1242/jcs.239681.

57. Galovic, M., Xu, D., Areces, L. B., van der Kammen, R. & Innocenti, M. Interplay between N-WASP and CK2 optimizes clathrin-mediated endocytosis of EGFR. Journal of Cell Science 124, 2001–2012 (2011).

58. Sigismund, S. et al. Clathrin-mediated internalization is essential for sustained EGFR signaling but dispensable for degradation. Dev Cell 15, 209–219 (2008).

59. Wary, K. K., Mariotti, A., Zurzolo, C. & Giancotti, F. G. A Requirement for Caveolin-1 and Associated Kinase Fyn in Integrin Signaling and Anchorage-Dependent Cell Growth. Cell 94, 625–634 (1998).

60. Elosegui-Artola, A. et al. Rigidity sensing and adaptation through regulation of integrin types. Nature Mater 13, 631–637 (2014).

61. Deyne, P. G. D. et al. The vitronectin receptor associates with clathrin-coated membrane domains via the cytoplasmic domain of its β5 subunit. Journal of Cell Science 111, 2729–2740 (1998).

62. Mrkonjić, S. et al. TRPV4 participates in the establishment of trailing adhesions and directional persistence of migrating cells. Pflügers Archiv - European Journal of Physiology 467, 2107–2119 (2015).

63. Franco, S. J. et al. Calpain-mediated proteolysis of talin regulates adhesion dynamics. Nat Cell Biol 6, 977–983 (2004).

64. Yan, B., Calderwood, D. A., Yaspan, B. & Ginsberg, M. H. Calpain Cleavage Promotes Talin Binding to the β3Integrin Cytoplasmic Domain. Journal of Biological Chemistry 276, 28164–28170 (2001).

65. Brodsky, F. M. Diversity of Clathrin Function: New Tricks for an Old Protein. Annu. Rev. Cell Dev. Biol. 28, 309–336 (2012).

66. Näthke, I., Hill, B. L., Parham, P. & Brodsky, F. M. The calcium-binding site of clathrin light chains. Journal of Biological Chemistry 265, 18621–18627 (1990).

67. Hazelbaker, M. A. et al. Arp2/3-mediated turnover of large clathrin lattices is regulated through the tyrosine kinase ACK. 2026.06.12.731920 Preprint at 10.64898/2026.06.12.731920 (2026).

68. Carrillo-Garcia, J. et al. The mechanosensitive Piezo1 channel controls endosome trafficking for an efficient cytokinetic abscission. Sci. Adv. 7, eabi7785 (2021).

69. Fielding, A. B., Willox, A. K., Okeke, E. & Royle, S. J. Clathrin-mediated endocytosis is inhibited during mitosis. Proceedings of the National Academy of Sciences 109, 6572–6577 (2012).

70. Kaur, S., Fielding, A. B., Gassner, G., Carter, N. J. & Royle, S. J. An unmet actin requirement explains the mitotic inhibition of clathrin-mediated endocytosis. eLife 3, e00829 (2014).

71. Eisenhoffer, G. T. et al. Crowding induces live cell extrusion to maintain homeostatic cell numbers in epithelia. Nature 484, 546–549 (2012).

72. Saitoh, S. et al. Rab5-regulated endocytosis plays a crucial role in apical extrusion of transformed cells. Proc Natl Acad Sci USA 114, E2327–E2336 (2017).

73. Lorenzo-Martín, L. F. et al. VAV2 signaling promotes regenerative proliferation in both cutaneous and head and neck squamous cell carcinoma. Nat Commun 11, 4788 (2020).

74. Panciera, T. et al. Reprogramming normal cells into tumour precursors requires ECM stiffness and oncogene-mediated changes of cell mechanical properties. Nat. Mater. 19, 797–806 (2020).

75. Martin, P., Pardo-Pastor, C., Jenkins, R. G. & Rosenblatt, J. Imperfect wound healing sets the stage for chronic diseases. Science 386, eadp2974 (2024).

76. Umesh, V., Rape, A. D., Ulrich, T. A. & Kumar, S. Microenvironmental Stiffness Enhances Glioma Cell Proliferation by Stimulating Epidermal Growth Factor Receptor Signaling. PLoS ONE 9, e101771 (2014).

77. Pardo-Pastor, C. & Rosenblatt, J. Piezo1 activates noncanonical EGFR endocytosis and signaling. Sci. Adv. 9, eadi1328 (2023).

78. Banushi, B., Joseph, S. R., Lum, B., Lee, J. J. & Simpson, F. Endocytosis in cancer and cancer therapy. Nat Rev Cancer 23, 450–473 (2023).

79. Scott, A. M., Wolchok, J. D. & Old, L. J. Antibody therapy of cancer. Nat Rev Cancer 12, 278–287 (2012).

80. Chew, H. Y. et al. Endocytosis Inhibition in Humans to Improve Responses to ADCC-Mediating Antibodies. Cell 180, 895–914.e27 (2020).

81. Thomas, R. & Weihua, Z. Rethink of EGFR in Cancer With Its Kinase Independent Function on Board. Front. Oncol. 9, 800 (2019).

82. Grove, J. et al. Flat clathrin lattices: stable features of the plasma membrane. MBoC 25, 3581–3594 (2014).

83. Wang, S. et al. Integrin αvβ5 Internalizes Zika Virus during Neural Stem Cells Infection and Provides a Promising Target for Antiviral Therapy. Cell Reports 30, 969–983.e4 (2020).

84. Marsh, M. & Helenius, A. Virus Entry: Open Sesame. Cell 124, 729–740 (2006).

85. Joseph, J. G. & Liu, A. P. Mechanical Regulation of Endocytosis: New Insights and Recent Advances. Adv. Biosys. 4, 1900278 (2020).

86. Li, J. H., Trivedi, V. & Diz-Muñoz, A. Understanding the interplay of membrane trafficking, cell surface mechanics, and stem cell differentiation. Seminars in Cell & Developmental Biology 133, 123–134 (2023).

87. Schindelin, J., et al. Fiji: an open-source platform for biological-image analysis. *Nat Methods* 9, 676–682 (2012).

88. McQuin, C. et al. CellProfiler 3.0: Next-generation image processing for biology. PLoS Biol 16, e2005970 (2018).

89. Vassilopoulos, S., Gibaud, S., Jimenez, A., Caillol, G. & Leterrier, C. Ultrastructure of the axonal periodic scaffold reveals a braid-like organization of actin rings. Nat Commun 10, 5803 (2019).

